# Systematic simulation of the interactions of Pleckstrin homology domains with membranes

**DOI:** 10.1101/2021.12.16.472954

**Authors:** Kyle I.P. Le Huray, He Wang, Frank Sobott, Antreas C. Kalli

## Abstract

Pleckstrin homology (PH) domains can recruit proteins to membranes by recognition of phosphatidylinositol phosphates (PIPs). Here we report the systematic simulation of the interactions of 100 mammalian PH domains with PIP containing model membranes. Comparison with crystal structures of PH domains bound to PIP analogues demonstrates that our method correctly identifies interactions at known canonical and non-canonical sites, while revealing additional functionally important sites for interaction not observed in the crystal structure, such as for P-Rex1 and Akt1. At the family level, we find that the β1 and β2 strands and their connecting loop constitute the primary PIP interaction site for the majority of PH domains, but we highlight interesting exceptional cases. Simultaneous interaction with multiple PIPs and clustering of PIPs induced by PH domain binding are also observed. Our findings support a general paradigm for PH domain membrane association involving multivalent interactions with anionic lipids.

**Teaser:** Simulating the binding of 100 Pleckstrin homology domains to cell membranes reveals patterns in their lipid interactions.

## Introduction

Peripheral membrane proteins (PMPs) are proteins which transiently associate with the surface of cellular or organelle membranes (*1, 2*). Binding of PMPs to membranes is often stabilized through a combination of specific and non-specific interactions with the lipid headgroups, insertion of hydrophobic regions of the protein into the membrane interior and/or the presence of post-translational modifications that can anchor the protein to the membrane. There are several families of structurally conserved protein domains whose members have been identified as membrane binding domains in PMPs. These families include C2 domains, Phox homology (PX) domains, FYVE domains, PDZ domains and Pleckstrin homology (PH) domains (*3*). Despite increasing structural and functional data about membrane binding domains, knowledge of their membrane binding interfaces, mechanism of association to the membrane and whether there is any common mechanism of association at the family-level, remain elusive. Some members of these families are capable of recognizing specific lipid species, such as phosphatidylinositol phosphates (PIPs). PIPs are a minority lipid component in membranes, but they play a substantial role in the regulation of membrane protein activity and cellular signaling (*4–6*).

PH domains are a large domain family with structural data available for more than 100 mammalian members. PH domains have a conserved fold (Fig. 1), which consists of a 7-stranded β-barrel, capped at one end by a C-terminal alpha helix, with a pocket at the open end that is typically positively charged for interaction with anionic PIP headgroups. The PH domain of PLCδ1 was the first identified domain capable of specific binding to PIP lipids (phosphatidylinositol-(4,5)-bisphosphate in particular), and membrane localization mediated via specific binding to PIPs is the most studied characteristic of PH domains (*7*). Some family members are known to participate in regulatory protein-protein interactions with other proteins, leading to the proposal that some PH domains do not have a membrane binding role (*8, 9*). However, the most recent literature indicates that the majority of the family members localize to membranes and interact specifically with phosphoinositides (*10, 11*).

**Fig. 1.**
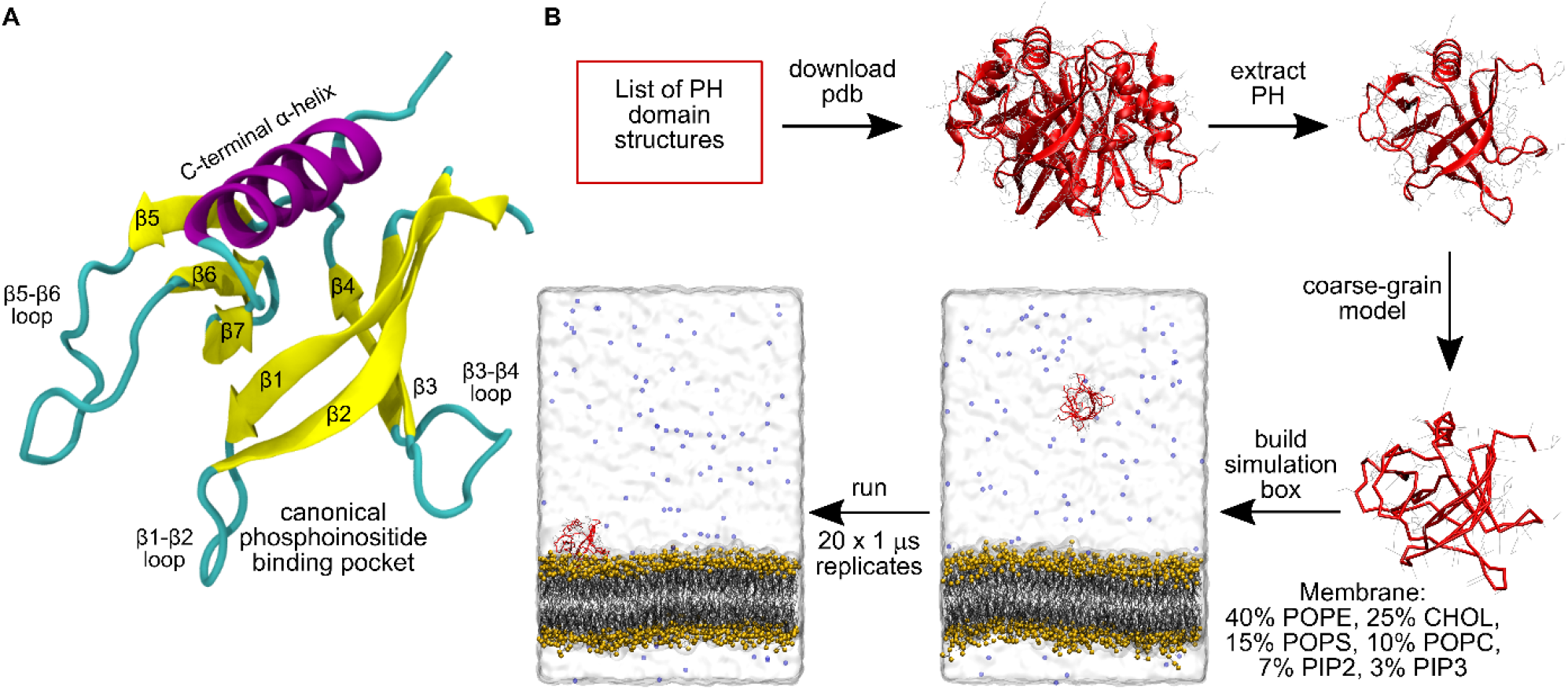
Conserved PH domain structure and simulation workflow. **(A)** Structure of the first PH domain of PLEK (PDB: 1xx0) demonstrates the conserved PH domain fold, consisting of a seven stranded β-barrel, capped by an α-helix, and with 6 variable inter-strand loops. The open end of the barrel contains the canonical pocket for phosphoinositide binding. **(B)** Illustration of the semi-automated simulation pipeline used for high throughput PH domain simulations in this study.

Due to their crucial role in the regulation and activity of many signaling proteins, PH domains are implicated in a number of diseases, including the Akt1, PDPK1, P-Rex1, and IQSEC1 PH domains in cancer and intellectual disability, the BTK PH domain in autoimmune disease and X-linked agammaglobulinemia, and the FGD1 PH domain in faciogenital dysplasia (*12–19*). Consequently, there is interest in the development of small molecule inhibitors of the PH domain-membrane interactions of these proteins, including recent work on inhibitors of P-Rex1 and IQSEC1 (*20–23*). Considering their importance in human disease and the pharmacological interest in mammalian PH domains, it is important to improve our understanding of their interactions with membranes. Most structures of PH domains that have been solved in complex with PIP lipid headgroups or suitable analogues typically demonstrate phosphoinositide binding at a so-called canonical site inside the pocket at the open end of the barrel (Fig. 1) (*14, 24–26*). In this pocket, the headgroup phosphates are stabilized by electrostatic and hydrogen-bonded interactions with the strands and unstructured loops flanking the cavity. The importance of basic residues in the loop region connecting the β1 and β2 strands (the β1-β2 loop; Fig. 1) for the interaction with PIPs has been shown, and a KX_n_(K/R)XR sequence motif in this region has been identified as a predictor of binding to PIPs phosphorylated at the 3 position (*27*). However, phosphoinositide binding at atypical sites on the exterior of the barrel (SI. Fig. 1) and by PH domains lacking this sequence motif has also been observed, for example in the case of the ArhGAP9 and β-spectrin PH domains (*28, 29*).

A structure of the ASAP1 PH domain revealed dual binding of anionic lipids to both canonical and alternate sites simultaneously (*30*). A combination of simulations and experiments later revealed that multiple anionic lipid binding maintains the ASAP1 PH domain in an orientation conducive for interaction with its membrane-bound protein target (*31*). Furthermore, recent evidence has been presented for three PIP interacting sites on the PLEKHA7 PH domain, and for PIP clustering induced by this PH domain (*32*). Similarly, an additional atypical site for soluble inositol hexakisphosphate binding has been observed in the BTK PH domain, which is critical for BTK activation (*33*). Subsequent simulations showed that multiple PIP binding sites stabilized dimerization of the BTK PH domain on the membrane (*34*). Additional sites for interaction with PIPs or anionic PS lipids have been identified in the Akt1, GRP1, PDK1 and BRAG2 PH domains (*35–40*). Furthermore, a large study of binding of yeast PH domains to liposomes of different composition demonstrated cooperative lipid binding in 93% of the liposome-binding PH domains (*10*). This growing body of evidence points to a new paradigm for PH domain membrane association, involving multivalent association with PIPs and other anionic lipids, rather than the one-to-one interaction mode suggested previously. It may be the case that the additional binding sites are weaker, more disordered or too dependent on the membrane environment to be resolved by crystallography. Despite these recent data, however, it remains unclear how widespread the capacity for multiple PIP binding is throughout the family of mammalian PH domains.

Molecular dynamics (MD) simulations of membrane protein structures computationally re-embedded into a lipid bilayer provide an excellent complement to experimental techniques and have proven to be a powerful tool for the identification of specific protein-lipid interaction sites (*41, 42*). In particular, previous simulations of the membrane interactions of 13 PH domains whose structures had been solved in complex with PIP headgroups or analogues found that such simulations can identify the crystallographic PIP binding sites, while also highlighting putative alternative sites of PIP interaction not revealed in the structures (*43*).

In this work, we simulated 100 mammalian PH domains, with the goal of establishing patterns in PH domain interactions with membranes. We find that the PH domain β1 and β2 strands and their connecting loop contains the primary contact site for PIP headgroups in 85% of the analyzed PH domains, and the majority of those have frequent contacts with PIPs at alternative sites, such as β3-β4 and β6-β7 regions. Our analysis highlights the diversity of PH domain membrane interactions and we have identified interesting exceptional cases. Furthermore, close association of multiple PIPs with the PH domains and clustering of PIPs induced by PH domain binding was universally observed in our simulations, with some PH domains exhibiting this to a greater extent than others.

## Results

### A semi-automated pipeline for coarse-grained molecular dynamics (CG-MD) simulation of PH domains

To perform simulations on this scale, we developed a semi-automated simulation pipeline (Fig. 1B). Given a PH domain containing structure, it will extract the PH domain, remodel any missing atoms or residues, and then build, energy minimize and equilibrate the simulation system. The PH domain is initially placed at a 6 nm z-axis distance from a symmetric lipid bilayer model composed of 10% POPC, 40% POPE, 15% POPS, 7% PIP_2_, 3% PIP_3_ and 25% cholesterol. For each PH domain 20 × 1 µs repeat simulations were conducted, each initialized with different velocities sampled from a Boltzmann distribution. We find that 20 replicates were sufficient for convergence in contacts analysis and protein-membrane distance analysis (Fig. S2). Not all simulation replicates resulted in stable membrane association, but membrane association was observed in the majority of replicates for all simulated PH domains (Fig. S3).

### Simulated phosphoinositide interaction sites are consistent with available crystal structures and suggest additional interaction sites

The capability of CG-MD simulations to identify crystallographic phosphoinositide binding sites on PH domains and other proteins has been previously demonstrated (*42, 43*). It is further demonstrated in our study for three PH domains with known crystallographic PIP binding sites that have not previously been simulated – the two PH domains of ADAP1 and the PREX1 PH domain, which are all known to possess canonical binding sites. Analysis of the number of contacts that each residue made with PIP_2_ and PIP_3_ headgroups during the final 200 ns of simulation and comparison with the relevant crystal structures (Fig. 2) shows that our simulations correctly predicted the binding of a phosphoinositide headgroup in the binding site suggested by the crystal structures for these three PH domains. We note that our contact analysis also identified additional interaction sites on the exterior of the β-barrel structure of these PH domains.

**Fig 2.**
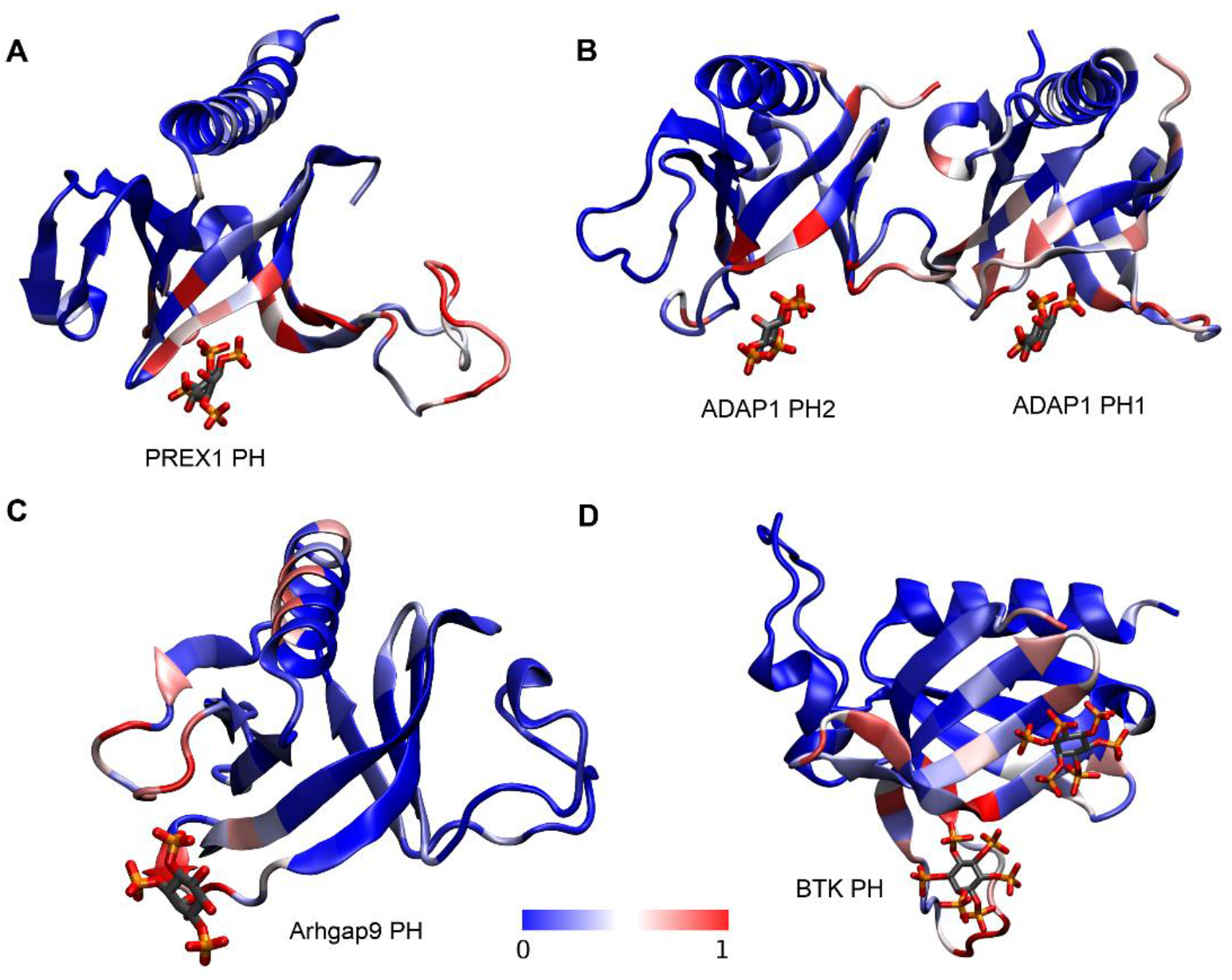
Comparison of simulated phosphoinositide interactions and crystallographic binding sites. Structures of the PH domains of **(A)** PREX1, **(B)** ADAP1, **(C)** Arhgap9, and **(D)** BTK in which each residue is colored according to the normalized number of contacts observed between the protein and PIP_2_ and PIP_3_ headgroups during the final 200 ns of simulation, averaged over 20 replicates. Normalization was carried out by dividing the number of contacts at every residue by the maximum number of contacts that any residue in that PH domain made with PIP headgroups. The position of the bound PIP-headgroup analogue in the PDB file of each structure is also shown (PDB IDs: 5D3X, 3LJU, 2P0H, 4Y94).

In the structure of the human PREX1 PH domain (PDB ID: 5D3X) in complex with inositol-(1,3,4,5)-tetrakis phosphate at the canonical site, examination of the side chains within 4 Å of the ligand shows that residues R289, R328, K368, K280 and Y300 are engaged in electrostatic and/or hydrogen-bonded interactions with the phosphates in the 3, 4 and 5 positions. S282 and Q287 also lie within 4 Å of the ligand and stabilize the interaction (*14*). During our simulations we find all these residues except Q287 have contacts with PIP headgroups at a frequency of at least 70% of that of the residue with the most contacts. Furthermore, K280 and R289 are the residues in the canonical binding site which make the most contacts with PIP headgroups in the simulations, which is consistent with experimental findings using differential scanning fluorimetry that these are the residues that are most important for PIP_3_ binding to the canonical pocket (*14*). In addition to binding at the canonical site, our analysis for the PREX1 PH domain also reveals substantial interaction of phosphoinositide headgroups with the disordered β3-β4 loop, which is rich in basic residues and has been modelled in for the simulations as it is absent from the structure. Experimentally, it has been shown that mutations which abolish binding to the canonical site do not abolish membrane binding in the PREX1 PH domain, but that membrane association is substantially reduced by deletion of β3-β4 loop residues 311–318 (*14*). This has led to a model of membrane interaction and activation in which non-specific electrostatic interaction between anionic lipids and the β3-β4 loop drive membrane association and thus allow PI(3,4,5)P_3_ binding to the canonical site, which then allosterically activates PREX1 (*14*). Our contacts analysis captures both of these key interaction sites.

Similarly, we observe regions of high PIP contacts defining the canonical binding pockets of both PH domains of ADAP1. In contrast, for the Arhgap9 PH domain, which lacks a canonical binding site, we do not observe these, but instead high numbers of contacts along outward facing residues of the β1 strand and β1-β2 loop, and the β5-β6 loop, defining the atypical binding pocket observed in the crystal structure. Additionally, we see high numbers of contacts along the face of the C-terminal helix which has a cluster of basic residues aligned along the putative membrane-binding interface. These non-specific interactions with anionic lipids potentially stabilize the membrane bound orientation of the protein. Lastly, for the BTK PH domain we observe high numbers of PIP contacts at both the canonical and atypical binding sites which were observed in the BTK PH/PIP headgroup crystal structure. Additional interactions along the β3-β4 loop, which may constitute a third interaction site. Overall, our contacts analysis captures both interaction sites, again demonstrating the power of our simulation method to reproduce crystallographic binding sites while adding detail of key interactions that are absent from some structures.

To examine these interactions in more detail for one PH domain, Akt1, the end point of one simulation in which PIP_3_ was bound to the known canonical site, was backmapped to an atomistic representation and simulated for a further 200 ns using the CHARMM36 force field (*25*). The interactions within the binding pocket (Fig. 3) are similar to the crystal structure, with R23, R25 and K39 engaged in electrostatic and hydrogen bonded interactions with the position 3 and position 4 phosphates. Additionally, we find that the hydrophobic tip of the β1-β2 loop formed by Y18 and I19 inserts into the membrane and engages in hydrophobic interactions with an acyl tail of the bound PIP_3_ and a cholesterol molecule. The β2-β3 and β3-β4 loops face away from the membrane, as has previously been suggested (*25*). In addition to interactions with the canonically bound PIP_3_, the membrane binding interface is lined with basic residues which facilitate association with 8 additional PIP_2_, PIP_3_ and PS lipids in the simulation snapshot (Fig. 3). In particular, there is a PIP_2_ bound in the pocket formed between the β1-β2 and β5-β6 loops (similar to the non-canonical binding site seen in the Arhgap9 PH domain), which is stabilized by interaction with R15, K20 and R67. Previous work has shown that R15 and K20 are critical for binding of the Akt1 PH to PS containing liposomes, and we propose that the other basic residues lining the membrane binding interface are likely to also contribute towards stabilization of the Akt1 PH domain on the membrane, involving multivalent interactions with anionic lipids (*35*).

**Fig. 3.**
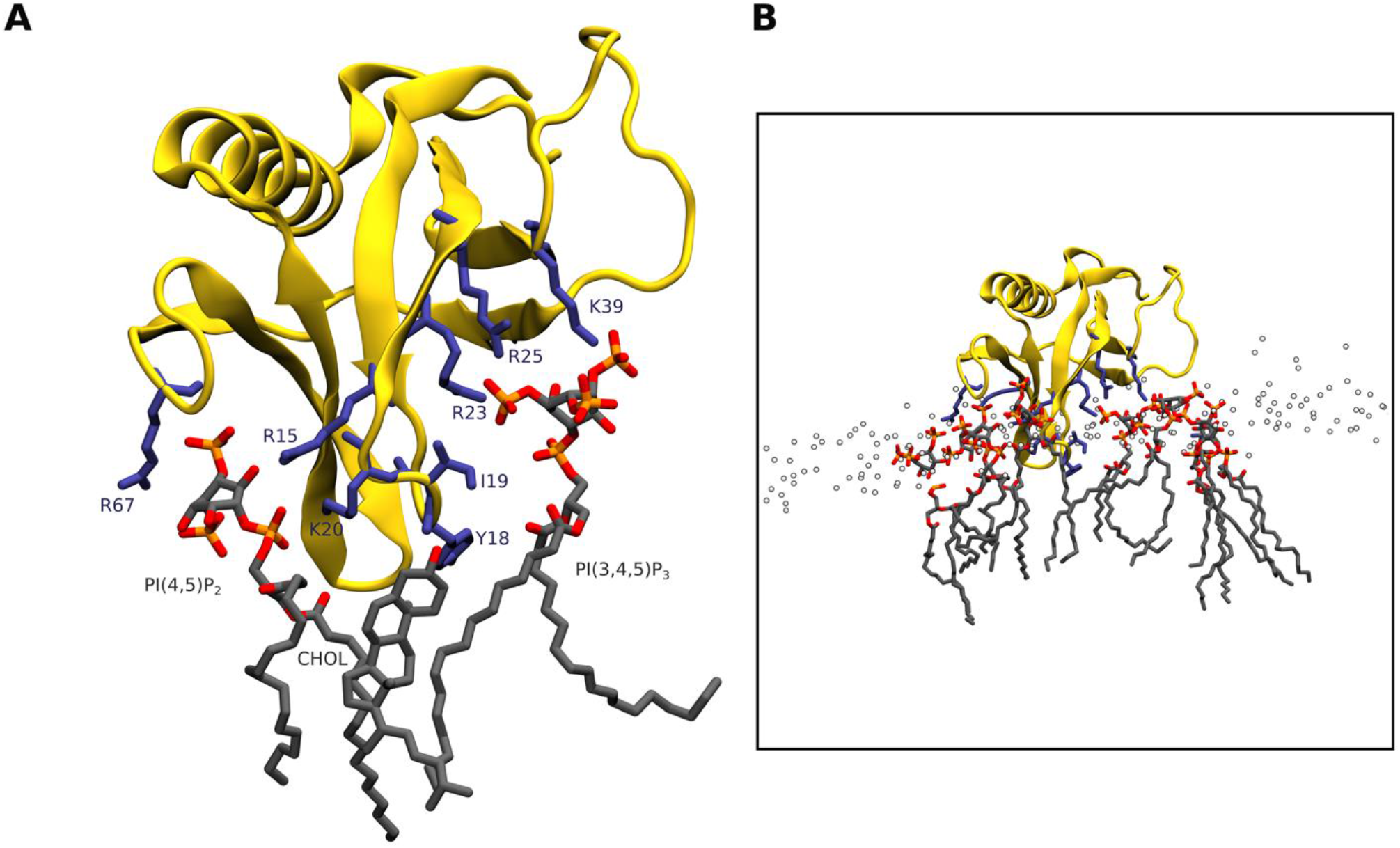
An atomistic model of membrane-bound Akt1 PH after backmapping and 200 ns of atomistic simulation. **(A)** Geometry of PI(3,4,5)P_3_ bound in the canonical site, with PI(4,5)P_2_ bound in a putative non-canonical site on the opposite face of the β1-β2 loop, meanwhile the hydrophobic tip of the loop inserts into the membrane and engages in hydrophobic interactions with cholesterol and lipid acyl tails. **(B)** the membrane bound state involves association of multiple anionic lipids with the PH domain.

### The β1-β2 region provides the primary site of PIP contacts in most PH domains

To obtain a global view of PIP-PH domain interactions, we examined the contribution to phosphoinositide headgroup contacts during simulations from each of the conserved secondary structure segments found in PH domains, allowing us to establish patterns and exceptions in the phosphoinositide contact profile across the family. Assigning the residues of the simulated PH domains to one of 14 secondary structure segments (7 strands, 6 interstrand/loop regions or the C-terminal helix) we totaled the PIP headgroup contacts of each segment during the final 200 ns of all simulations. To determine the frequency of contacts at each segment, we normalized to also take into consideration the sequence length of the segments. For the purpose of this analysis, the short, structured regions sometimes found between the classical PH domain strands have been assigned to the loop between the strands. This analysis (Fig. 4A) reveals that the β1-β2 loop is the segment most likely to interact with PIPs, followed by the β3-β4 loop, β2 strand and β6-β7 loops. The importance of the β1-β2 loop has long been known, but our study also showed significant interactions of the β3-β4 loop and β6-β7 loops. These loops form a triad at the base of the beta sheets in PH domains with canonical binding sites, but the very high number of contacts of β3-β4 loop and β6-β7 loops also suggests association of multiple PIP lipids with the PH domains during the simulations. A similar analysis can be applied at the amino acid level, highlighting the importance of cationic lysine, arginine and histidine residues for stabilizing PIP headgroup interactions (Fig. 4B).

**Fig. 4.**
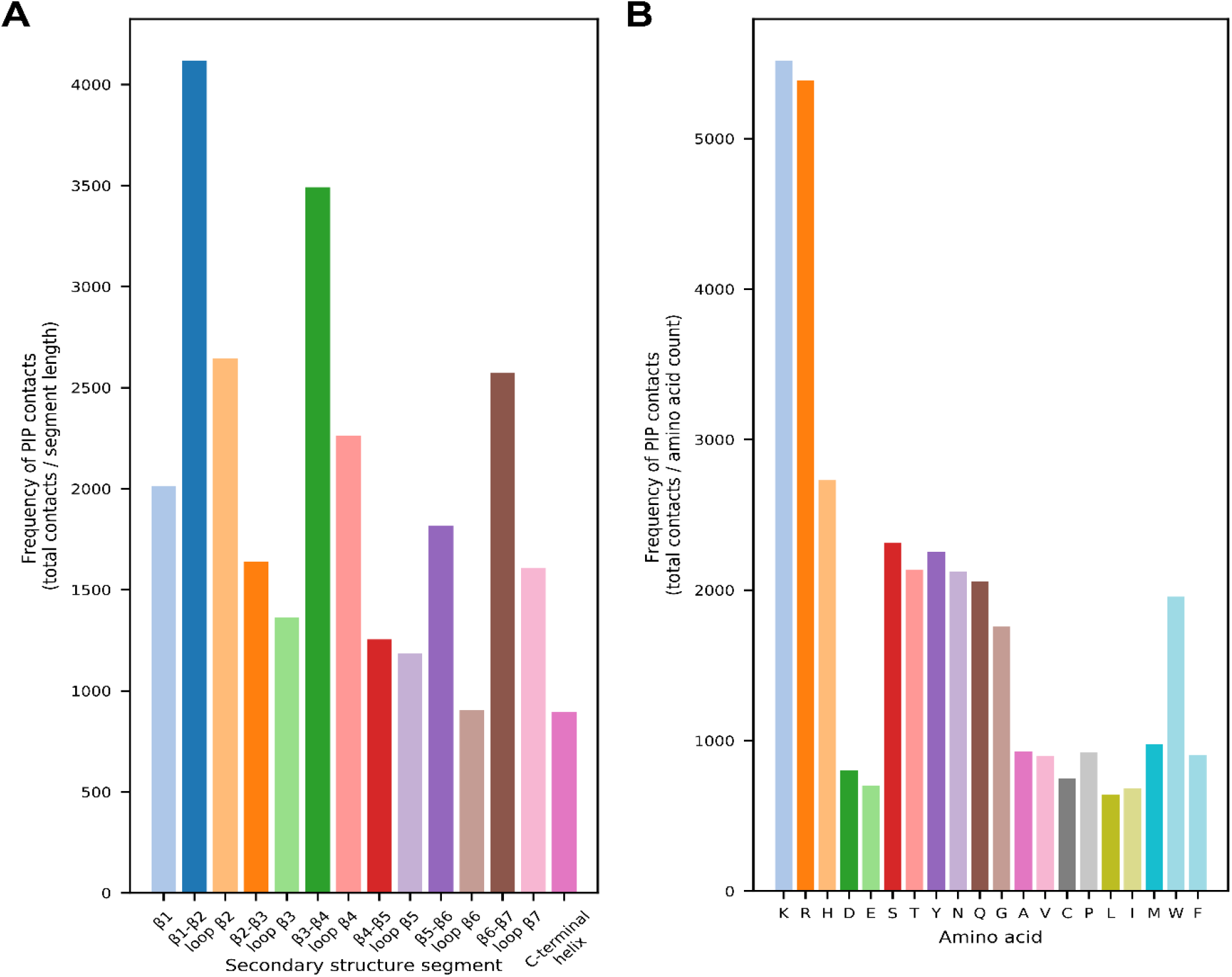
Secondary structure and amino acid contributions to phosphoinositide contacts. **(A)** Frequency with which each of the secondary segments of PH domains made contacts with phosphoinositide headgroups during all simulations. PH domain residues were assigned to one of 14 conserved secondary structure units and contact frequency was calculated by summing contacts for each residue assigned to the secondary structure unit over all PH domains and simulation replicates and dividing by the total number of residues assigned to that secondary structure unit. **(B)** frequency with which each amino acid type contacted PIP_2_ and PIP_3_ headgroups during all simulations. Frequency was calculated by summing contacts for the amino acid over all simulation replicates of all PH domains and dividing by the total number of occurrences of that amino acid in the simulated sequences.

We next examined the contact frequency of secondary structure segments for individual PH domains. Residue-level contacts with PIP_2_ and PIP_3_ headgroups were totaled during the final 200 ns of simulation for each PH domain, and normalized by dividing the maximum number of contacts made by a single residue in that PH domain. To reduce the complexity of the analysis, we selected a normalized contact frequency of 0.8 as a threshold for a residue with substantial contributions towards PIP headgroup interactions in the PH domain, as this threshold captured the key residues for known crystal binding sites. Using this threshold, we determined whether each secondary structure segment contains a residue which contributes substantially to PIP interactions (Fig. 5)

**Fig. 5.**
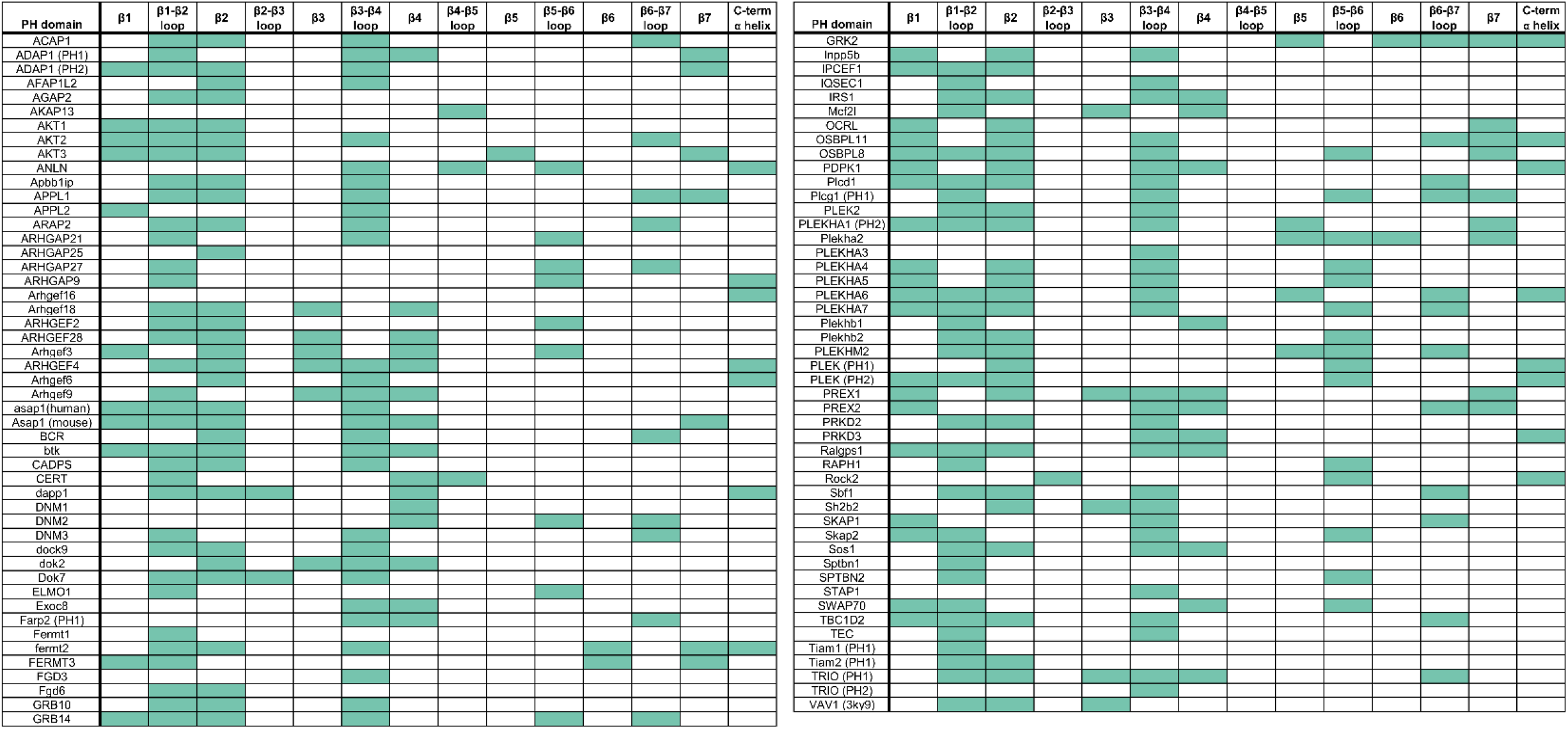
Identification of structural segments with substantial contributions towards PIP interactions for individual PH domains. Summary of simulated PH domain contacts with phosphoinositide headgroups, identifying whether or not each of the 14 conserved structural elements of the PH domain had a residue with normalized contacts above a threshold value of 0.8. Those with contacts in a segment above the threshold are colored green, otherwise white. Some PH domains that couldn’t be reasonably assigned to the classical PH domain secondary structure pattern were omitted for this analysis (see methods).

Using this analysis, we found that 85% of the analyzed PH domains (including those lacking the canonical KX_n_(K/R)XR motif in this region) possess substantial PIP interactions at a residue in β1, β2, or the connecting β1-β2 loop. Consistent with our global analysis, this indicates functional conservation of the β1-β2 region for PIP binding. Furthermore, among those PH domains with the primary contact site in the β1-β2 region, 89% of those have additional contacts of similar frequency at alternative sites such as β3-β4 and β6-β7, pointing to the supplementary role of these loops in stabilizing the primary PIP binding site and/or in interacting with additional PIPs.

This systematic analysis of the location of PIP contact sites also reveals interesting exceptional cases that do not conform to the interaction patterns discussed above. The Exoc8 PH domain is one example, in which the β3-β4 loop is the primary contact site observed for interactions with phosphoinositide headgroups (Fig. 6). This PH domain lacks basic residues in the β1-β2 loop, which are key for electrostatic interaction with anionic PIP headgroups. Instead, the primary site for PIP interaction is formed by a pair of arginines at the tip of the β3-β4 loop. These electrostatic interactions are supplemented by insertion of hydrophobic residues spread along the membrane interacting interface from β2 to β4. Examination of the electrostatic profile of the Exoc8 PH domain reveals a long electropositive ridge on one side of the β-barrel, arising from β3, β4 and their connecting loop. This electropositive ridge is not mirrored on the opposite β1-β2 side of the barrel, and we find that the Exoc8 PH domain stably adopts a ‘side-on’ membrane-bound orientation, maximizing the contact between the positive ridge and the negative membrane surface. Exoc8 has recently been found to bind specifically to PI(4,5)P_2_, although there is currently no experimental insight into its mechanism of membrane association (*11*).

**Fig. 6.**
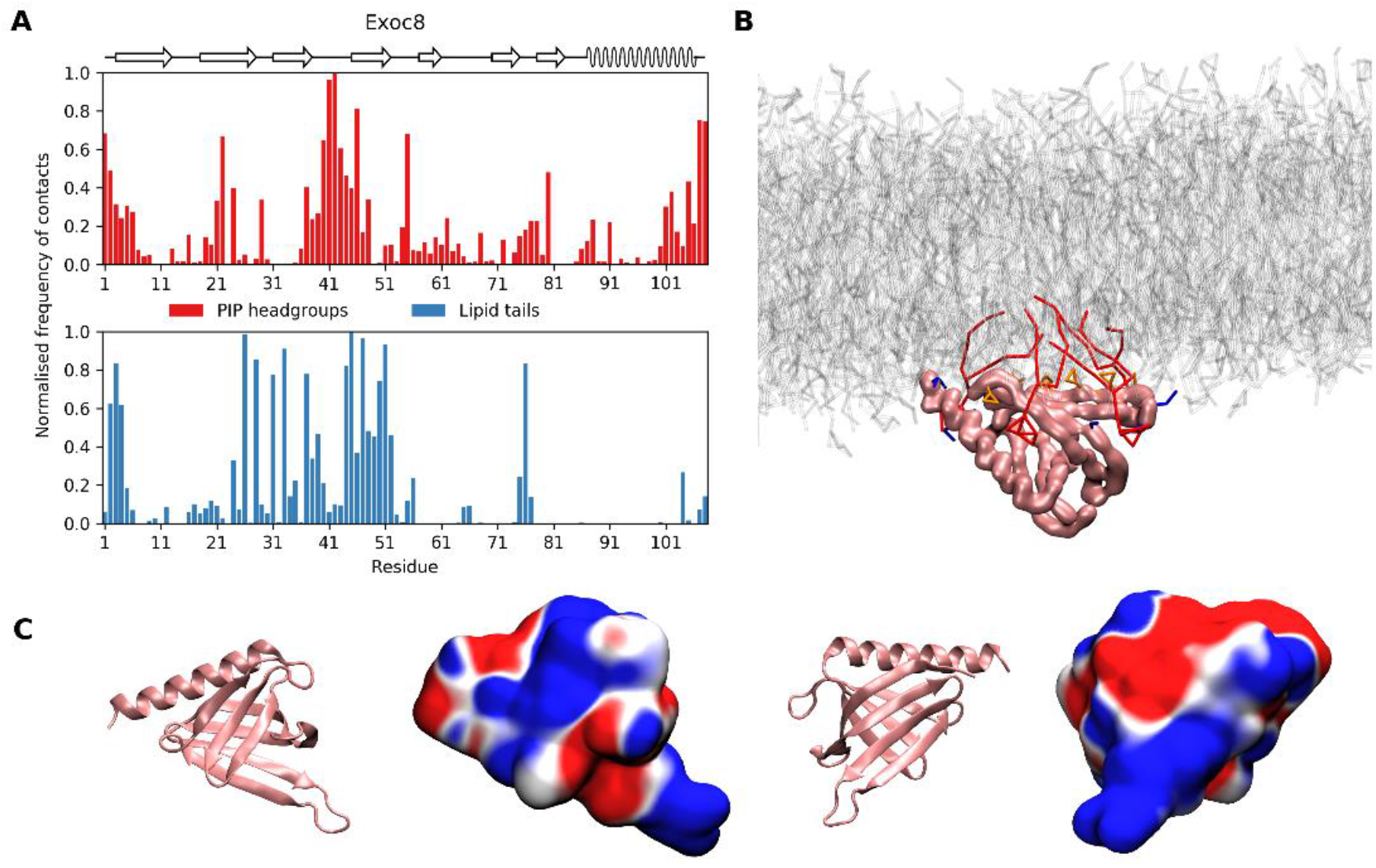
Interactions of the Exoc8 PH domain with the membrane. **(A)** Normalized number of contacts between the Exoc8 PH domain and PIP headgroups (red) or lipid tails (blue) reveals a preference for phosphoinositide interaction with the β3-β4 loop and not the β1-β2 loop. **(B)** Snapshot showing the preferred membrane bound orientation of Exoc8 PH. The lipid bilayer is shown in grey, with the PIP lipids that associate with the Exoc 8 PH domain shown in red. **(C)** Structure and electrostatic potential map of Exoc8 PH in two orientations, demonstrating the electropositive β3-β4 region.

### Association of multiple PIPs with PH domains

For all simulated PH domains, we observed that after initial binding to the bilayer, multiple PIPs are recruited and closely associate with the PH domain (Fig. 7). In the final 200 ns of simulation (using 0.65 nm cutoff distance to the PO4 phosphate particle), most PH domains have at least 4 PIPs within the cutoff distance (Fig. S4). Furthermore, clustering of PIP lipids in the vicinity of PH domains induces modifications to the local lipid environment, as seen through analysis of the lipid radial distribution function during the final 200 ns of simulation (Fig. S5). Other recent computational studies which have examined multiple phosphoinositide binding to PMPs have employed total phosphoinositide compositions ranging from 5-10% (*31, 32, 34, 43–45*). Our model membrane has concentrations of PIP_2_ (7%) and PIP_3_ (3%) which are at the upper end of this range. This PIP rich model may bias the simulation towards multiple PIP binding. To test this possibility, we repeated the simulations at lower PIP concentrations (3% PIP_2_, 1% PIP_3_) for the Plcδ1 PH domain. Similar multiple PIP association with the PH domain was observed at this lower PIP concentration after 2 µs of simulation (Fig. S6). This shows that whilst clustering may take longer with a different composition, the phosphoinositide concentration is not biasing our observation of phosphoinositide clustering.

**Fig. 7.**
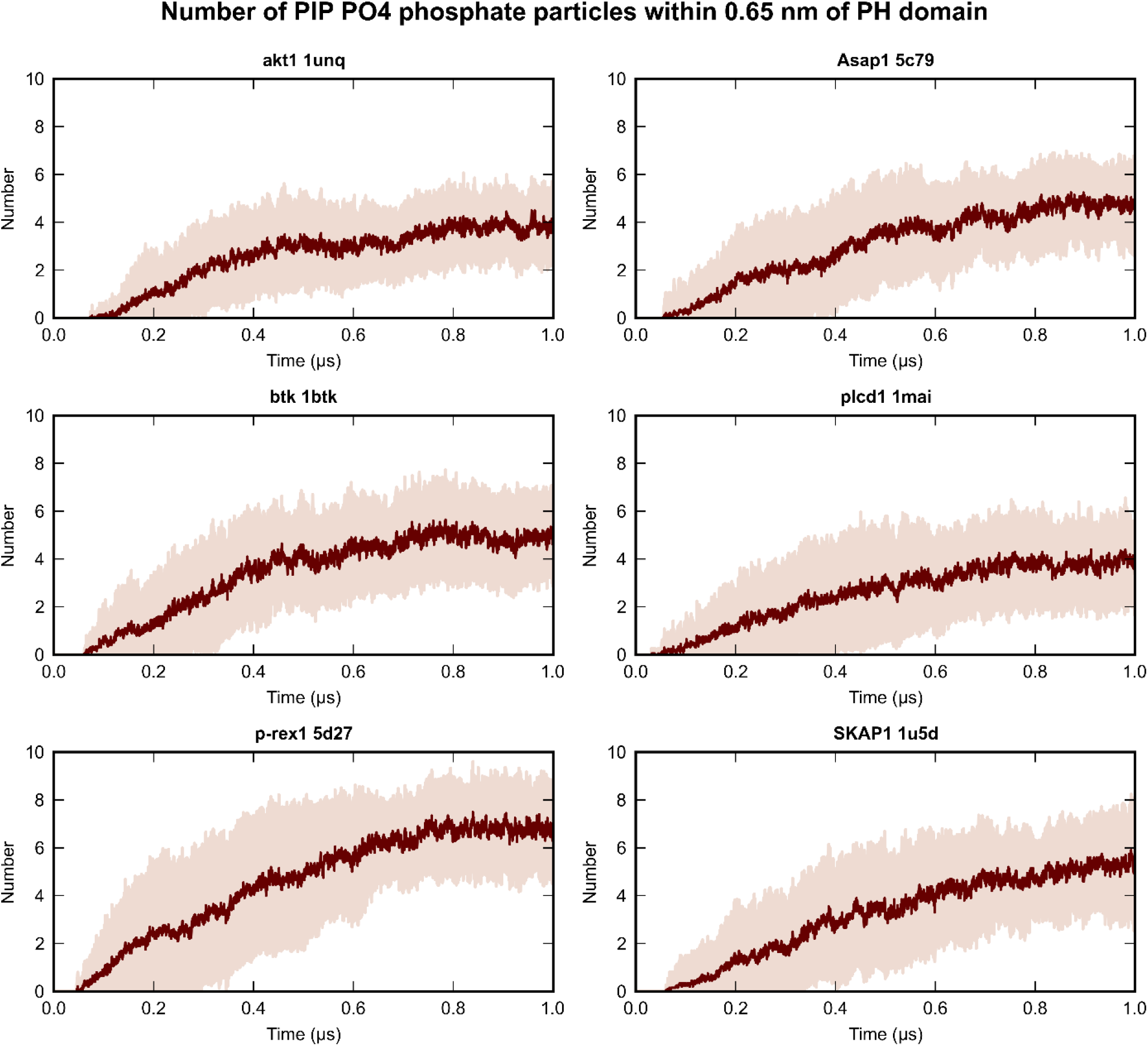
Multiple phosphoinositide molecules associate with PH domains during simulation. Plots of the number of PO4 (CG representation of position 1 phosphate) particles of PIP_2_ and PIP_3_ lipids within a 0.65 nm cutoff distance of the PH domain during simulations. The mean of 20 simulations for the AKT1, Asap1, btk, plcd1, p-rex1 and SKAP1 PH domains is plotted in dark red, with pale red shading representing the interval +/−1 standard deviation of the mean. The data for all simulated PH domains are shown in Fig. S4.

### PH domains adopt diverse membrane orientations

Lastly, we examined different preferred orientations of PH domains on the membrane. The PH domains of BTK (PDB: 1btk) and CYTH2 (PDB: 1u29) have crystal structures with inositol phosphate binding at the canonical site, whereas binding to the atypical binding site between the β1-β2 and β5-β6 loops is seen in the crystal structure of ArhGAP9 (PDB: 2p0h). These PH domains adopted different preferred membrane bound orientations during our simulations, dictated by differences in electrostatics. BTK preferentially associated in an orientation in which all loops except β5-β6 are in contact with the membrane, and the canonical PIP binding site is occupied. The β5-β6 loop in this PH domain is rich in glutamic acid residues and points away from the membrane surface due to electrostatic repulsion (Fig. 8A). In contrast to BTK, the PH domain of arhgap9 adopts a side-on orientation, with the positively charged face of the barrel containing the β1-β2, β4-β5 and β5-β6 loops in contact with the membrane. The β2-β3 and β3-β4 loops point away from the membrane. This orientation of the arhgap9 PH domain enables phosphoinositide headgroup binding to the atypical site formed between the β1-β2 and β5-β6 loops. The PH domain of Cyth2 adopts an orientation that is intermediate between that of BTK and arhgap9. It does not have the negatively charged β5-β6 loop that maintains the BTK PH domain in an ‘upright’ orientation. Its electrostatic potential map is asymmetric around the barrel, leading to a side-on orientation, but it is not quite as asymmetric as arhgap9. Interestingly, we observe phosphoinositide contacts with the β5-β6 loop in Cyth2, suggesting a non-canonical PIP binding site that is not observed in the crystal structure. However, the normalized frequency of contacts in this region is less than in arhgap9. This comparison suggests that electrostatics are important in determining the interaction with anionic lipids and the orientation of the PH domain on the membrane.

**Fig. 8.**
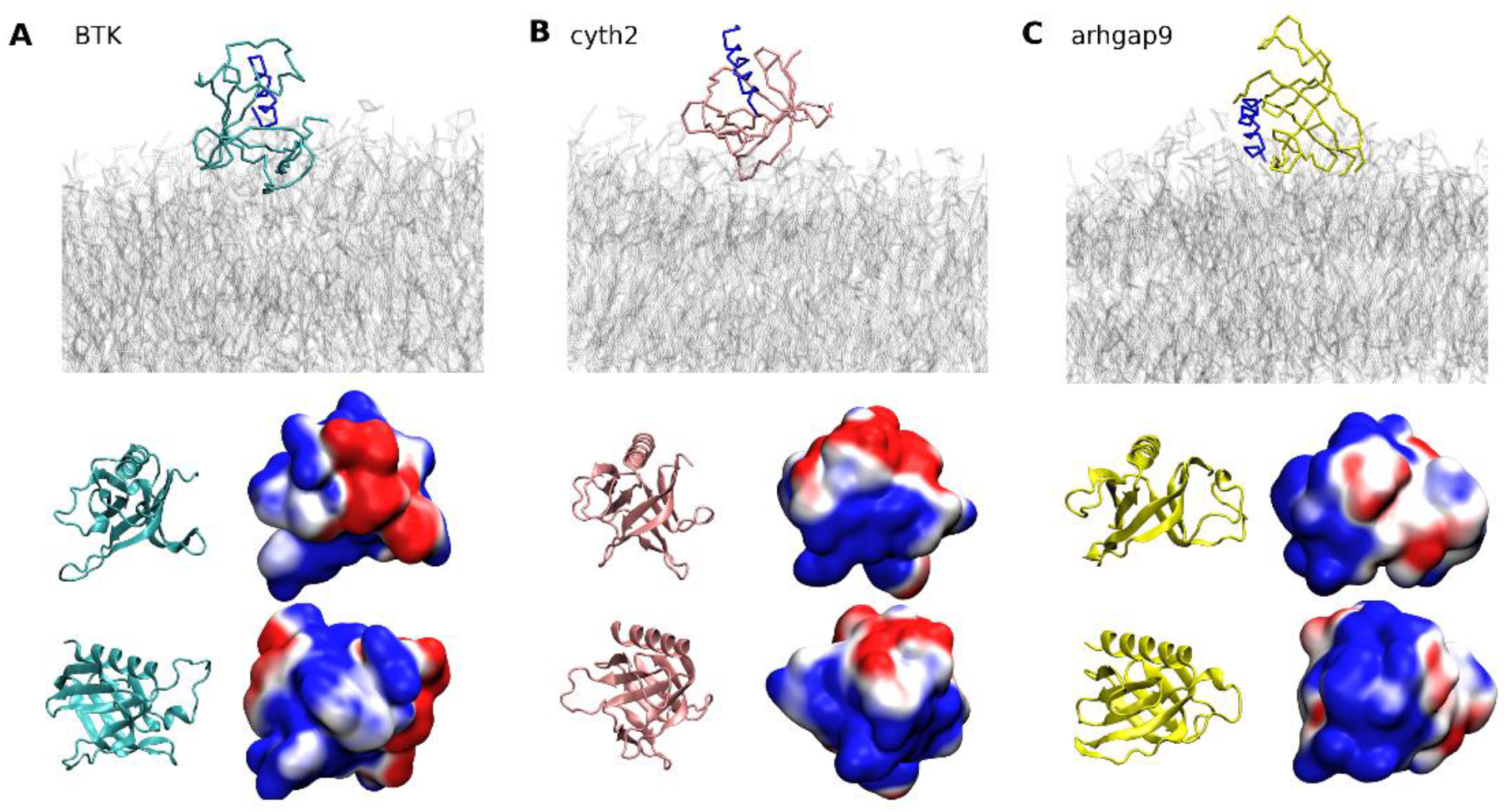
Diversity of preferred membrane bound orientations of PH domains. Snapshot of the preferred membrane bound orientation of btk **(A)**, cyth2 **(B)**, and arhgap9 **(C)** PH domains, with the C-terminal helix shown in blue to highlight the orientational differences. Further details of PIP headgroup and lipid tail contacts and orientational landscape are presented in SI. Fig. 7, with the orientations shown here selected from the most dense state in the orientational landscape shown in SI. Fig. 7.

## Discussion

Using high-throughput CG-MD simulations we have systematically compared the membrane interactions of 100 mammalian PH domains. We have observed that the β1-β2 region in PH domains is the primary site which makes contacts with PIPs in the majority of family members, but in most cases is supplemented by interactions at other sites, particularly the β3-β4 and β5-β6 loops. The significance of the β1-β2 loop for phosphoinositide binding has been suggested previously for a number of PH domains in both canonical and atypical binding modes, supplemented by adjacent loops – the β3-β4 loop in the canonical binding mode and the β5-β6 loop in the arhgap9-like atypical binding site (*12, 14, 46, 47*). Other regions have also been shown to play a role in phosphoinositide atypical binding in some PH domains, such as the β3 and β4 strands in the BTK PH domain, and the β6-β7 loop in the CERK PH domain (*33, 34, 48*).

Following initial membrane binding, additional PIPs are recruited, eventually saturating between 4-6 PIPs closely associating (< 0.65 nm) with the typical PH domain. This multiple PIP association occurs even using membrane compositions with lower (4%) total phosphoinositide concentrations. There is a growing body of evidence that membrane association by individual PH domains involves interaction with multiple PIPs, background anionic lipids such as POPS, or other lipid types such as sphingolipids (*10, 14, 30–32, 37, 49, 50*). Multiple binding sites for PIPs or soluble phosphoinositides have been identified for the ASAP1, BTK, PLEKHA7, dynamin and acap1 PH domains (*30, 32–34, 50, 51*). Enhancement of membrane binding by the presence of background anionic lipids such as POPS has been observed for the AKT1 and ASAP1 PH domains (*31, 35*). Beyond PH domains, recent studies have highlighted the role of multiple PIP binding sites in the membrane association of Phox homology (PX) domains, the AENTH complex involved in clathrin-mediated endocytosis and the TUBBY domain (*45, 52, 53*). These multiple interactions have been proposed to influence the orientation, localization, affinity and diffusivity of membrane associating proteins, and may be regulated by fluctuations and gradients in anionic lipid concentrations in different membranes (*31, 54–56*).

Our systematic simulations suggest that capacity for interaction with multiple PIPs, and PH domain induced lipid clustering is a general property of membrane associating mammalian PH domains, consistent with large scale studies of cooperative lipid binding in yeast PH domains (*10*). However, the relative strength and specificity of these multiple interaction sites remains to be determined. Local lipid modulation has been previously observed in simulations of integral membrane proteins, for which the altered local lipid environment provides a unique fingerprint (*57*). Our simulations suggest a similar behavior for PH domains. The PH domains may create a unique fingerprint in the membrane enriched with anionic lipids, which may regulate their interactions with partner proteins. Alternatively, recognition of already existing PIP clusters in the membrane could localize them to particular membrane regions or to the vicinity of an integral membrane protein interaction target.

It is important to consider some limitations of our methodology. We have used coarse-grained molecular dynamics simulations which use some approximations in modelling of both the protein and the lipids. Despite these approximations, the consistency of our findings with much of the recent literature on individual PH domains suggests that these simulations have the capacity to identify rather accurately the regions of important lipid interactions, including those that have not been captured in structural studies; although recent experiments are revealing some of these additional interactions, they remain difficult to identify at the molecular level systematically (*31, 32*). Additionally, our simulation method does not adequately sample membrane unbinding events that would enable us to determine the thermodynamics and relative affinities of PH domain binding to our model membrane system. The preferred binding orientations in our simulations can, however, provide a starting point for the use of biased simulation methods, such as the generation of a potential-of-mean-force using umbrella sampling, that would allow calculation of the strength of the PH domain-membrane association (*58*). Furthermore, although the coarse-grained model captures the preference that PH domains have for interaction with PIPs over other lipid types, the resolution of the model is not sufficient to accurately capture the experimental specificity that some PH domains display for PIP phosphorylation levels or positional isomers, hence our analysis focuses on PIP_2_/PIP_3_ interaction in general. However, PH domain specificities and binding affinities have been extensively studied and experimental techniques enable these to be characterized systematically (*11, 59*).

Understanding the protein-lipid interactome is crucial to improve our understanding of membrane protein structure, function and pharmacology, but it remains largely uncharacterized due to the diversity and complexity of membrane lipids and limitations in experimental techniques (*41, 60*). The increasing realism and throughput of coarse-grained molecular dynamic simulations is poised to allow systematic characterization at the molecular level. There are a few examples in which the lipid interactions of multiple proteins in complex bilayers have been systematically simulated in a single study (*43, 61–63*). Furthermore, the MemProtMD database contains simulations of over 5000 integral membrane structures in a single component DPPC lipid bilayer (*64*). The present work, covering 100 proteins and 2 ms of aggregate simulation time demonstrates the growing power of high-throughput simulations to systematically study protein-lipid interactions and to investigate the patterns and differences in lipid binding within families. Our approach can be readily extended to other membrane binding domain families, such as PX domains and C2 domains.

## Materials and Methods

### Discovery, selection and processing of PH domain structures for simulation

Uniprot advanced websearches were conducted and cross-referenced with the Protein Data Bank to find all reviewed PH domain containing sequences, with available PDB structures for human, rat or mouse. This produced a list of 115 PH domains. The unusual ‘split’ and ‘BEACH-type’ PH domains of, for example, NBEA, Plcg1 and Snta1 were subsequently excluded as unsuitable for simulation and comparison with other PH domains. PH domains were chosen for simulation ad hoc. Where a protein contained two distinct PH domains both were simulated but only one structure was simulated per PH domain. Structures were selected on the basis of resolution, low number of missing residues and the absence of mutations. Selected PDBs were downloaded and processed for simulation. Processing involved extracting the PH domain from the rest of the structure and truncating 2-4 residues before the first PH domain β-strand and 2-4 residues after the C-terminal α-helix to have a consistent structure for all the PH domains simulated. MODELLER was used to restore unresolved atoms or residues to the PH domain structure and to mutate any residues deviant from the wild-type uniprot sequence (*65*). Typically, only a few missing residues needed to be remodelled in the unstructured loop regions. Electrostatic potential maps of PH domain structures were generated using the PDB2PQR and APBS tools, at pH 7 with the CHARMM forcefield (*66*).

### CG-MD simulations

Simulations were performed using GROMACS 5.0.7 and the Martini 2.1 forcefield (*67, 68*). An automated script was developed and used to quickly and consistently build and equilibrate the simulation system and generate run files for production simulations. Processed PH domain PDB structures were converted to a CG representation using the martinize tool provided by the Martini developers and placed in a 16.5 nm × 16.5 nm × 20.5 nm simulation pbc box (*68*). The insane tool was used to add ions (0.1 mol L^-1^ Na^+^ and Cl^-^) and solvent water and construct a symmetric membrane bilayer (both leaflets composed of 10% POPC, 40% POPE, 15% POPS, 7% PIP2, 3% PIP3 and 25% cholesterol) at a z distance of 7-8 nm from the protein (*69*). An elastic network model with a 0.7 nm cutoff distance was applied to protein backbone particles to constrain secondary and tertiary structure (*70*). All systems were energy minimized and then equilibrated in the NPT ensemble for 2 ns with protein backbone particles restrained. For each system 20 production simulations were run for 1 µs, with each repeat simulation initialized with random velocities according to a Maxwell-Boltzmann distribution. The LINCS algorithm was used to constrain bonds to equilibrium length (*71*). The velocity rescaling method was used to maintain a temperature of 323 K, with a 1 ps coupling time. Semi-isotropic Parrinello-Rahman coupling was used to maintain a pressure of 1 bar using a 12 ps coupling time (*72, 73*).

### Atomistic MD simulations

The final frame of the 19^th^ replicate simulation of the Akt1 PH domain was selected for backmapping to an all-atom representation due to the presence of PIP_3_ in the known canonical binding pocket. Backmapping of the system from Martini to the CHARMM36 force field was achieved using the backward method and the initram.sh script provided by the Martini developers (*74*). To correct for any structural changes within the protein during the CG simulation and backmapping, the backmapped protein coordinates were replaced with those from the original crystal structure (PDB ID: 1unq) after superimposition of the 1unq structure upon the backmapped structure using the confrms GROMACS command. CHARMMGUI provides parameters for several different PIP_2_ and PIP_3_ isomers – here we used POPI25 and POPI35, which have a proton on the position-5 phosphate, on the basis of ab initio calculations indicating that this is the most stable protonation state of PI(4,5)P_2_ and the evidence that Akt1 does not form strong interactions with the position 5 phosphate of inositol tetraphosphate (*25, 75*). To ensure the correct headgroup stereochemistry of the PIP_3_ bound at the canonical site after backmapping, the headgroup coordinates of this POPI35 were replaced (after superimposition by rmsconf) by those of a reference POPI35 obtained from a pure POPI35 membrane constructed using the CHARMM-GUI membrane builder (*76*). The backmapped system was subsequently energy minimized and subject to 1 ns of equilibration in the NPT ensemble with the protein backbone restrained. An unrestrained production simulation was run for 200 ns, with a 2 fs timestep, temperature of 323 K and semi-isotropic Parrinello-Rahman pressure coupling at 1 bar.

### CG-MD analysis

The protein was first centered in the trajectory using gmx trjconv to prevent artifacts in analysis arising due to periodic boundary conditions. Root-mean-square deviation (gmx rms) and root-mean-square fluctuation (gmx rmsd) of protein backbone particles were calculated for each trajectory relative to its first frame. Z-axis distance between protein and membrane centres were calculated using the gmx dist command. Analysis of protein orientation was achieved by finding the rotation matrix (gmx rotmat) describing the transformation between a reference orientation and the orientation at each frame. The reference orientation was arbitrarily selected as the orientation in the end frame of the first simulation in the set of replicates. A Python script was developed for generating orientation density plots. These were constructed by taking the z-dist and the R_zz_ component of the rotmat data, correcting the rotmat to account for the membrane symmetry (R_zz_ value at a frame is multiplied by −1 if the z-dist at that frame is negative), then generating and plotting a 2D histogram in matplotlib.

Contacts between all residues and lipids were calculated for the following lipid groups: POPC headgroup, POPE headgroup, POPS headgroup, POP2 headgroup, POP3 headgroup, all CHOL particles, POP2 and POP3 headgroups combined and all lipid tails. gmx mindist was used to calculate whether a molecule possessing particles belonging to the given lipid group were within a 5.5 Å cut-off distance of any protein particles belonging to each residue at each simulation frame during the final 200 ns. One contact was counted at the residue for every frame in which a lipid particle in the group was within the cutoff distance. For each PH domain and each lipid group, contact counts at each residue were totaled across the 20 repeat simulations, and then normalized by dividing the total contacts at each residue by the total contacts made by the residue with the highest number of contacts with that lipid group. This gives the normalized number of contacts, in which the residue with the most contacts has the value 1 and all other residues have the contacts normalized relative to this residue.

Convergence analysis was carried out as above, using differently sized samples of simulation replicates.

To calculate the number of PIP lipids closely associated with the PH domain during the time course of the simulation, gmx mindist was used to calculate the number of PO4 particles (the connecting phosphate between the tail and headgroup) belonging to either POP2 or POP3 lipids within a 0.65 nm distance of any protein particles. The mean and standard deviation was calculated over 20 simulation replicates for each PH domain. A larger cutoff distance was used here than for the contacts analysis due to the extra distance between the headgroup phosphates and the connecting PO4 phosphate. Radial distribution functions were similarly calculated using gmx rdf and the PO4 particles of each individual lipid species, using the final 200 ns of simulation from all 20 replicates.

### Family-wide comparison of contacts by amino acid or secondary structure

For the family-wide analysis of amino acid contacts with PIP headgroups shown in Fig. 4, we grouped all residues by their amino acid type and totaled the contacts counts (for the POP2 and POP3 headgroup lipid group) across all PH domains and simulation replicates. Normalization was conducted by dividing the total contacts for each amino acid type by the number of incidences of that amino acid in all the simulated PH domain sequences. The secondary structure analysis was plotted similarly, but with the contacts instead grouped according to the conserved PH domain structural segments. An in-house python script was developed for this grouping, which uses the STRIDE secondary structure assignment program to assign PH domain residues to one of 14 secondary structure segments given their structure (*77*). Normalization of the contacts was conducted by dividing the total contacts of each segment by the number of residues assigned to that segment over all the PH domains.

The 14 secondary structure segments were: β1 strand, β1-β2 loop, β2 strand, β2-β3 loop, β3 strand, β3-β4 loop, β4 strand, β4-β5 loop, β5 strand, β5-β6 loop, β6 strand, β6-β7 loop, β7 strand, and the C-terminal α-helix. Unstructured n-terminal residues before the first strand were assigned to β1 strand and unstructured C-terminal residues were assigned to the C-terminal α-helix. Several PH domains have short, structured regions inserted between the conserved structural segments, which complicates this analysis; to handle these cases we assigned the additional short helices or strands to the nearest loop region where reasonable through manual correction of the STRIDE file. Similarly, due to inaccuracies with STRIDE or structure resolution there are gaps in the strands of some PH domains and these were also corrected by this analysis where reasonable by manually assigning the appropriate residues to the correct strand in the STRIDE file. The PH domains for which the STRIDE file was manually edited for this analysis were: akap13 (6bca), anln (2y7b), ARHGAP27 (3pp2), arhgap9 (2p0h), Arhgef18 (6bcb), ARHGEF2_5efx, Arhgef3 (2z0q), Arhgef6 (1v61), Arhgef9 (2dfk), DNM3 (5a3f), FERMT3 (2ys3), inpp5b (2kig), PLEK PH1 (1pls), p-rex1 (5d27), PREX2 (6bnm), sptbn1 (1btn). Three PH domains with additional structured regions that did not reasonably fit in with this analysis were excluded from this part of the analysis, these PH domains were: ARHGEF1 (3odo), cyth2 (1u29) and cyth3 (1u29).

### Contacts threshold table

The residues of each PH domain (except ARHGEF, cyth2, and cyth3) were assigned to one of the 14 conserved structural segments as described above. For each segment in the PH domain, we used a python script to determine (TRUE/FALSE) whether or not that segment contained a residue with normalized frequency of contacts with POP2+POP3 headgroups above 0.8 (in other words a residue with total contacts equal to at least 80% of the residue that had the most contacts in that PH domain). This gives a family-wide overview of which segments of each PH domain provide the key contribution to PIP interaction.

## Supplementary Materials

**Fig. S1.**
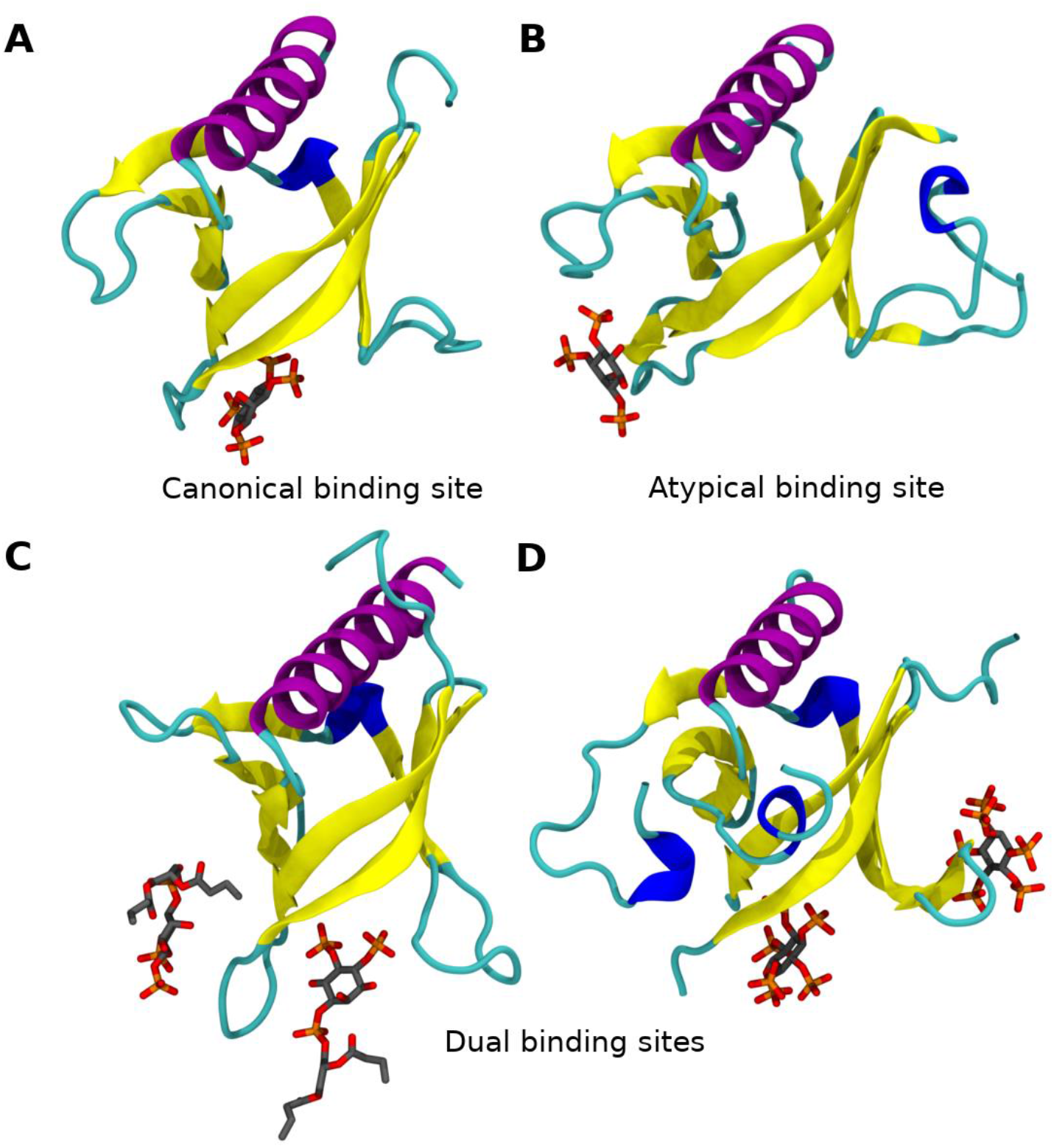
Example canonical and atypical phosphoinositide binding sites observed in PH domain crystal structures. **(A)** Structure of the DAPP1 PH domain (PDB: 1fao), demonstrating inositol tetraphosphate bound at the canonical site. **(B)** Structure of the ArhGAP9 PH domain (PDB: 2p0h), with inositol trisphosphate bound at an atypical site on the outside of the barrel. **(C)** Structure of the Asap1 PH domain (PDB: 5c79), exhibiting dual binding of 04:0 PI(4,5)P_2_ at both the canonical and an atypical site simultaneously. **(D)** Structure of the PH domain of BTK (PDB: 4y94), exhibiting dual binding of inositol hexa-kis-phosphate to both the canonical site and an atypical site distinct from that of Asap1 or ArhGAP9.

**Fig. S2.**
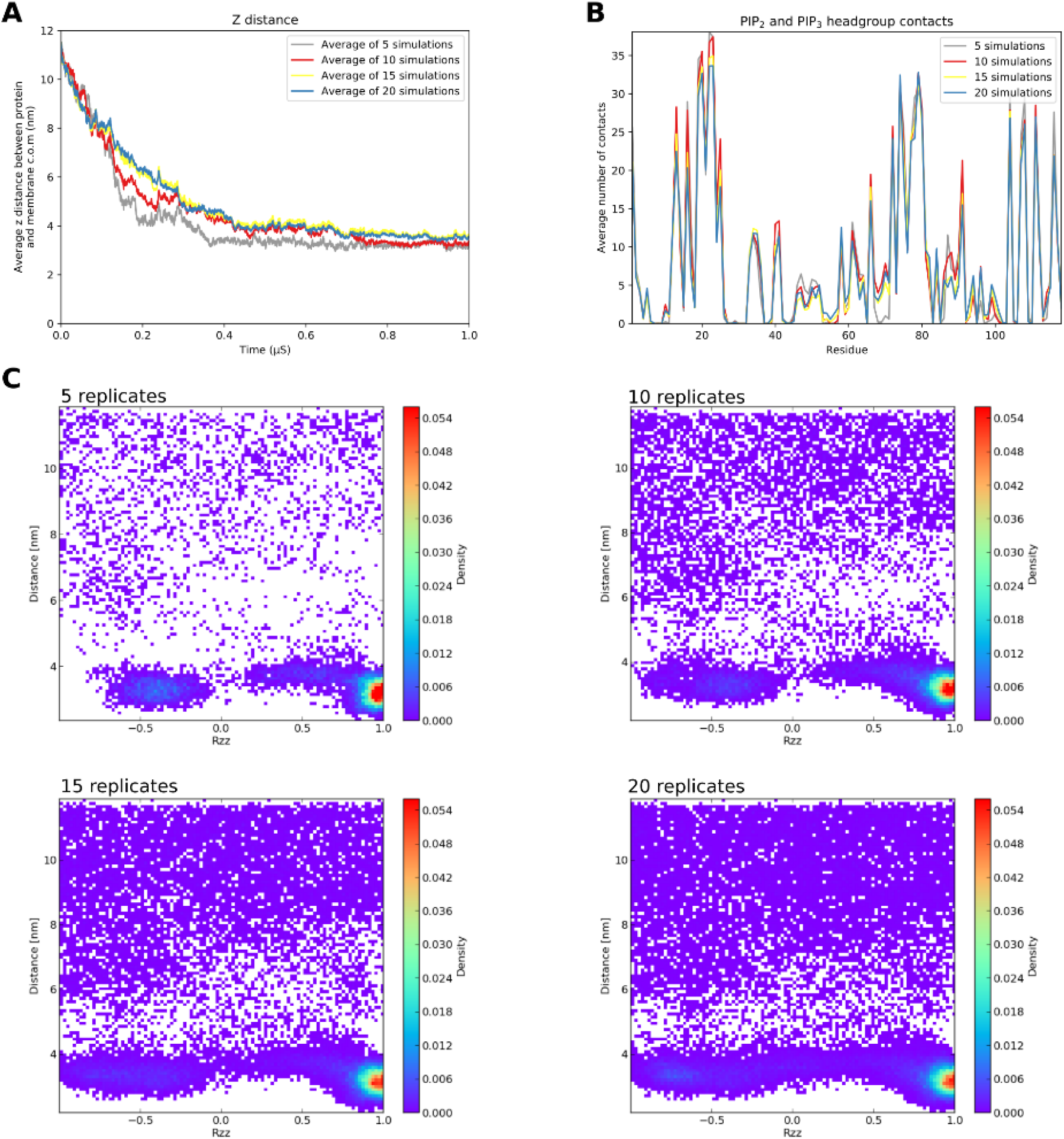
Convergence of simulation data. Analysis of simulations for the BTK PH domain, using data from 5, 10, 15 and 20 replicates. 20 simulation replicates achieves convergence in: **(A)** protein-membrane z-axis distance over time, **(B)** contacts with PIP_2_ and PIP_3_ headgroups and **(C)** distance-rotation density analysis.

**Fig. S3.**
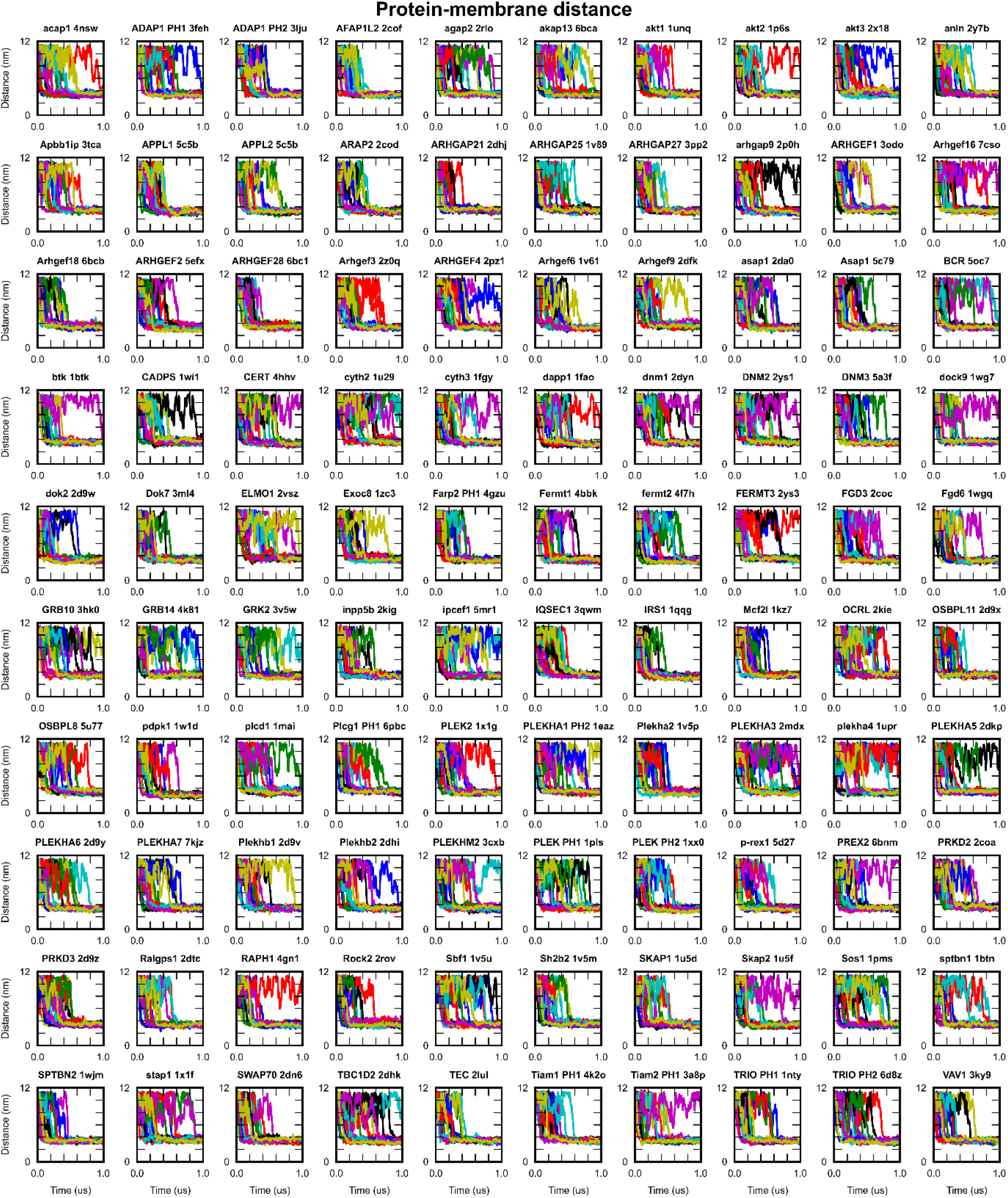
Z-axis distance between protein and membrane centres of mass during the simulation time course. Data shown for all PH domain simulations, where each color represents the trajectory for an independent replicate. Membrane binding is observed at a distance of approximately 4 nm. Distances have been corrected to account for periodic boundary conditions.

**Fig. S4.**
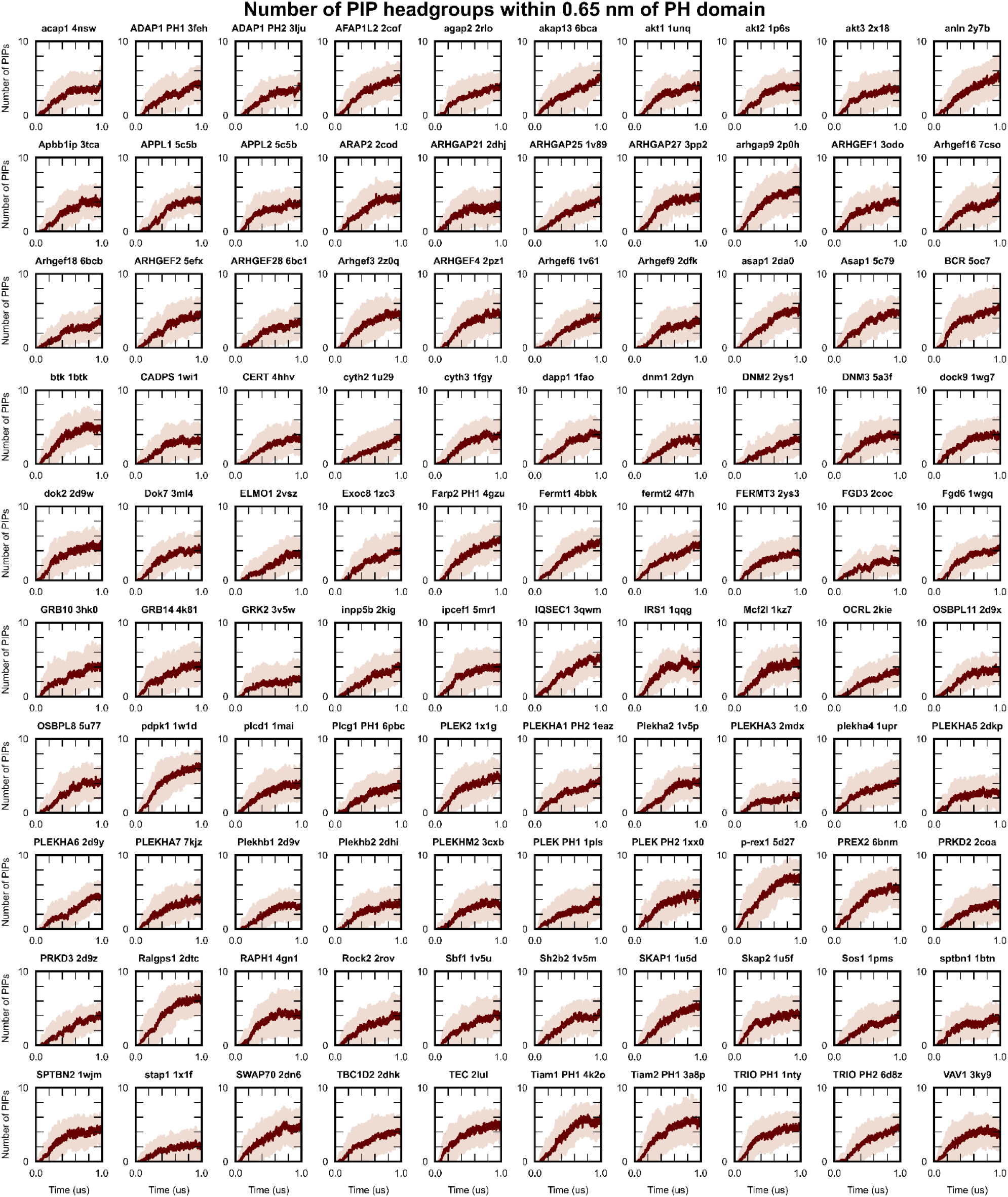
Association of multiple PIP lipids with PH domains following membrane binding. The number of PO4 (CG representation of position 1 phosphate) particles of PIP_2_ and PIP_3_ lipids within a 0.65 nm cutoff distance of each PH domain during the course of simulations. The mean of 20 simulations is plotted in dark red, with pale red shading representing the interval +/−1 standard deviation of the mean for every PH domain.

**Fig. S5.**
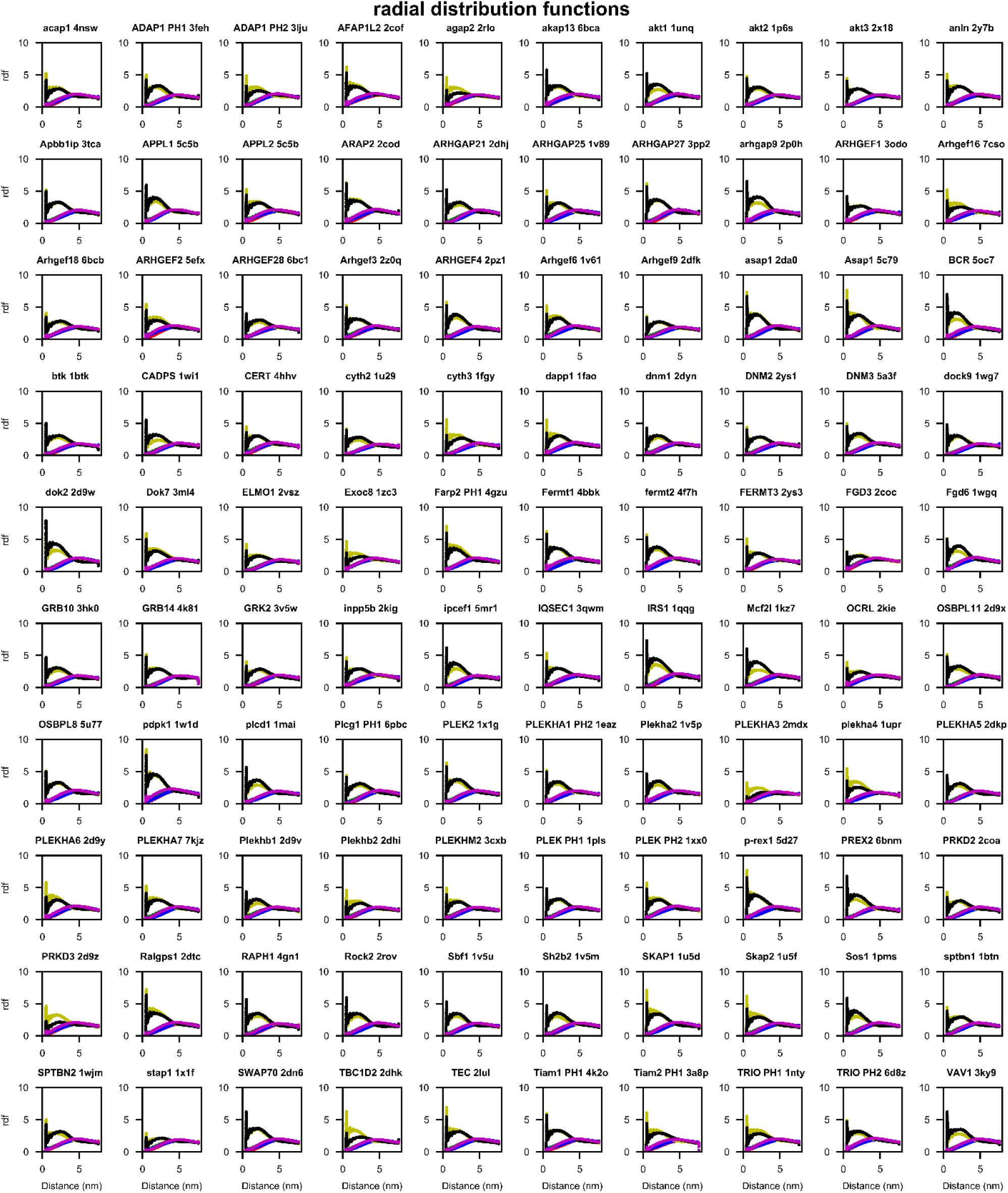
Lipid radial distribution functions around each PH domain demonstrate clustering of PIP lipids. Radial distribution functions around the protein were calculated using gmx rdf and the PO4 particles of each individual lipid species (PIP_2_: yellow, PIP_3_: black, cholesterol: magenta, POPS: blue, POPE: green, POPC: red) during the final 200 ns of simulation of all replicates for each PH domain.

**Fig. S6.**
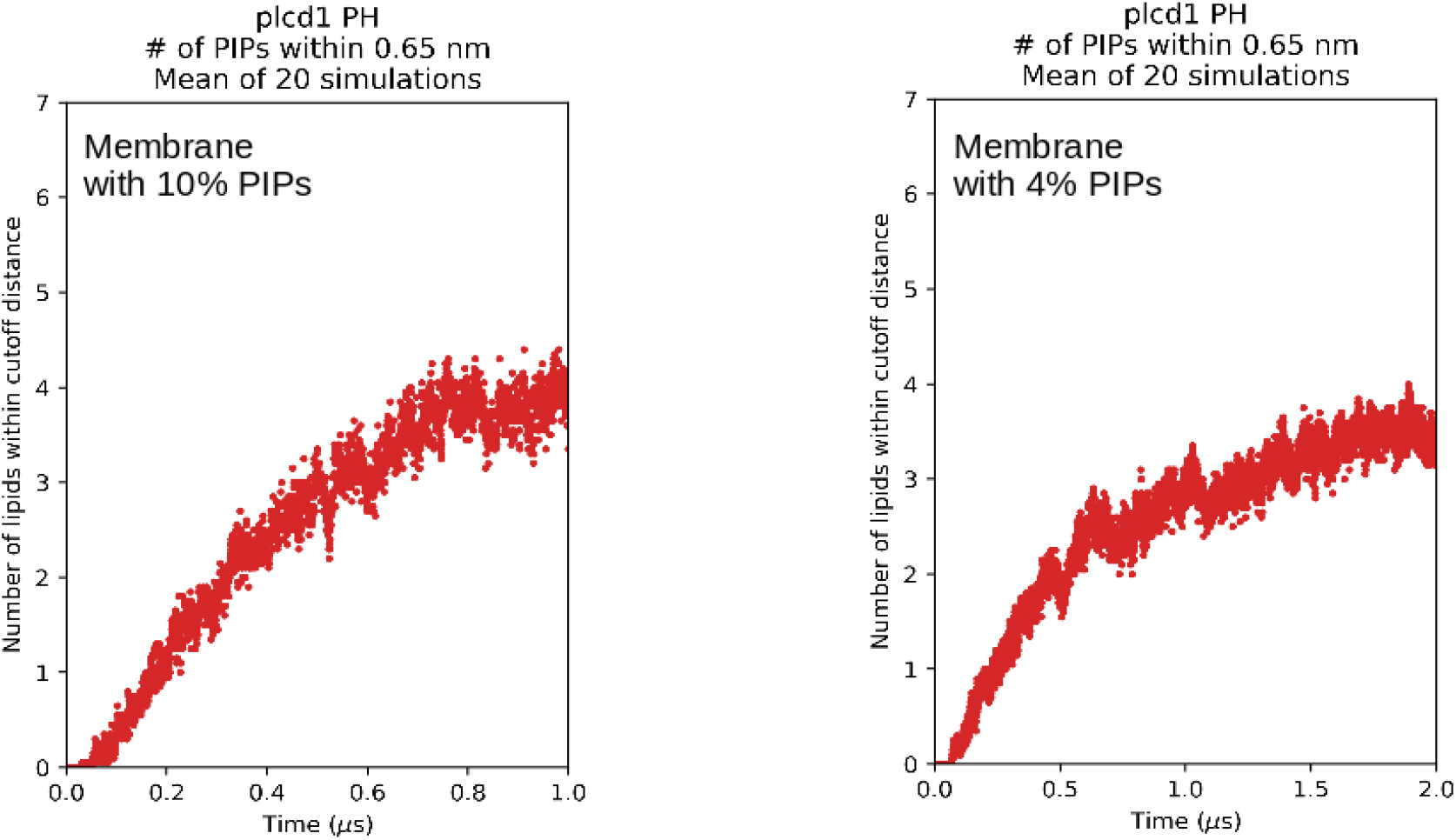
Association of multiple PIPs with the plcd1 PH domain in membranes with 10% and 4% PIP composition. The number of PO4 (CG representation of position 1 phosphate) particles of PIP_2_ and PIP_3_ lipids within a 0.65 nm cutoff distance of the PH domain during simulations. The mean of 20 simulations is plotted in red.

**Fig. S7.**
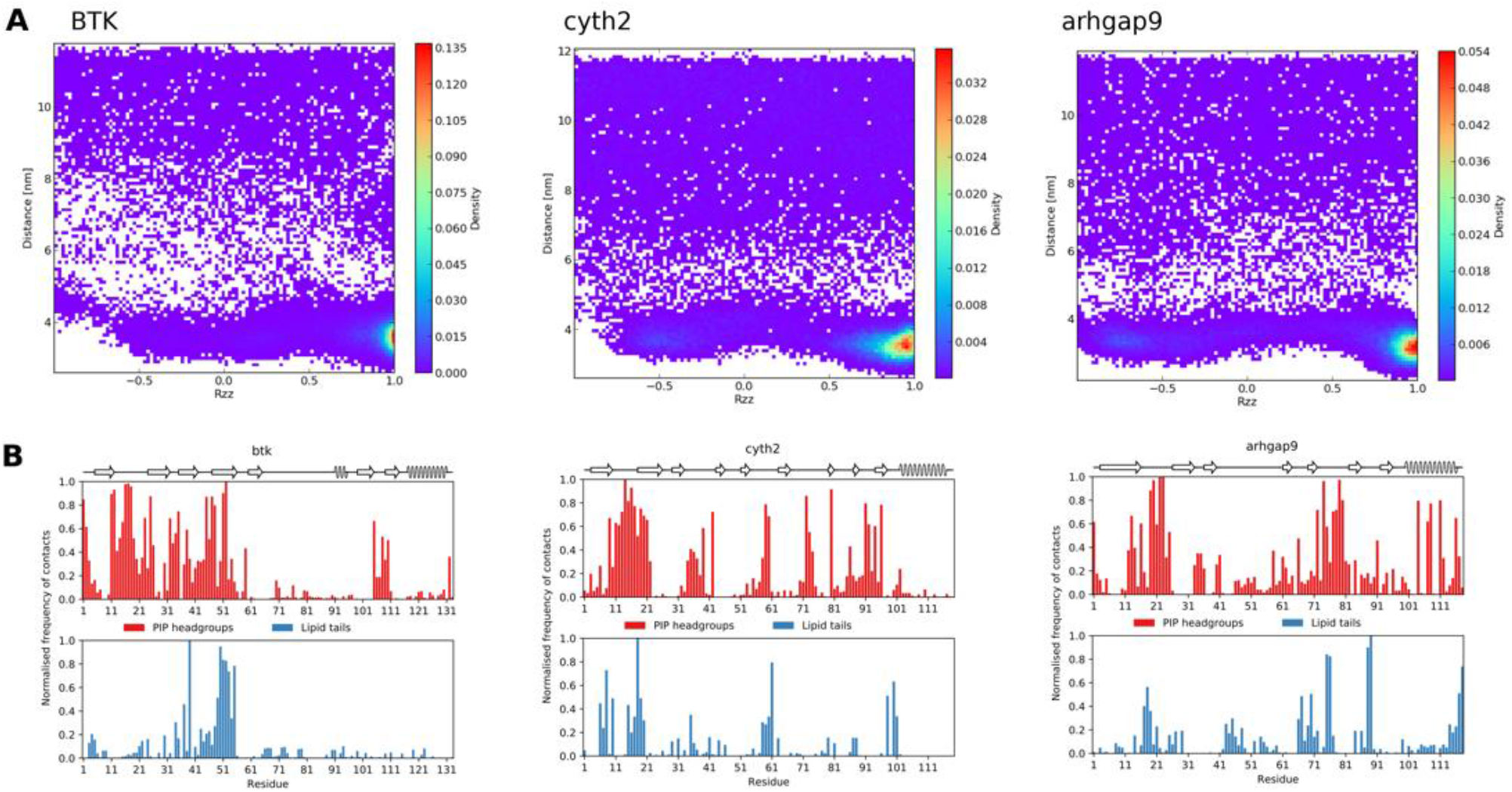
Supplement to Fig. 7. **(A)** rotation-distance density matrices, and **(B)** normalized contacts between the protein and PIP_2_ and PIP_3_ headgroups (red), or phospholipid tails (blue) for the BTK, cyth2, arhgap9 PH domains.

**Fig. S8.**
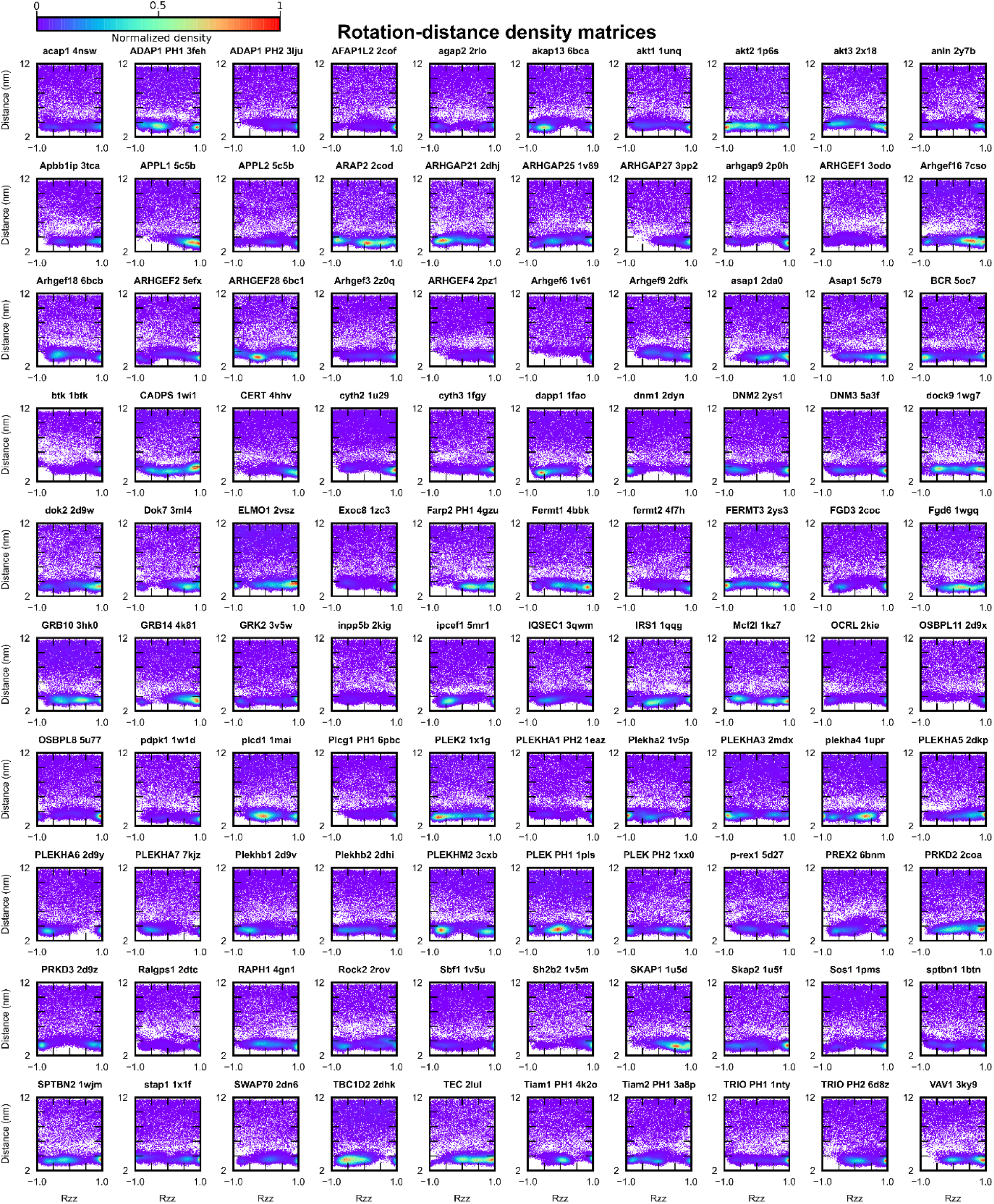
PH domains adopt preferred orientations on the membrane, as demonstrated by rotation-distance density matrices. Each 2D histogram show the density of protein-membrane z-distance and orientation states observed during simulation for on PH domain. The orientation was characterized by the Rzz component of the rotation matrix describing the transformation between an arbitrary reference orientation and the PH domain at each frame, corrected for periodic boundary conditions. Reference orientations differ for each PH domain.

**Fig. S9.**
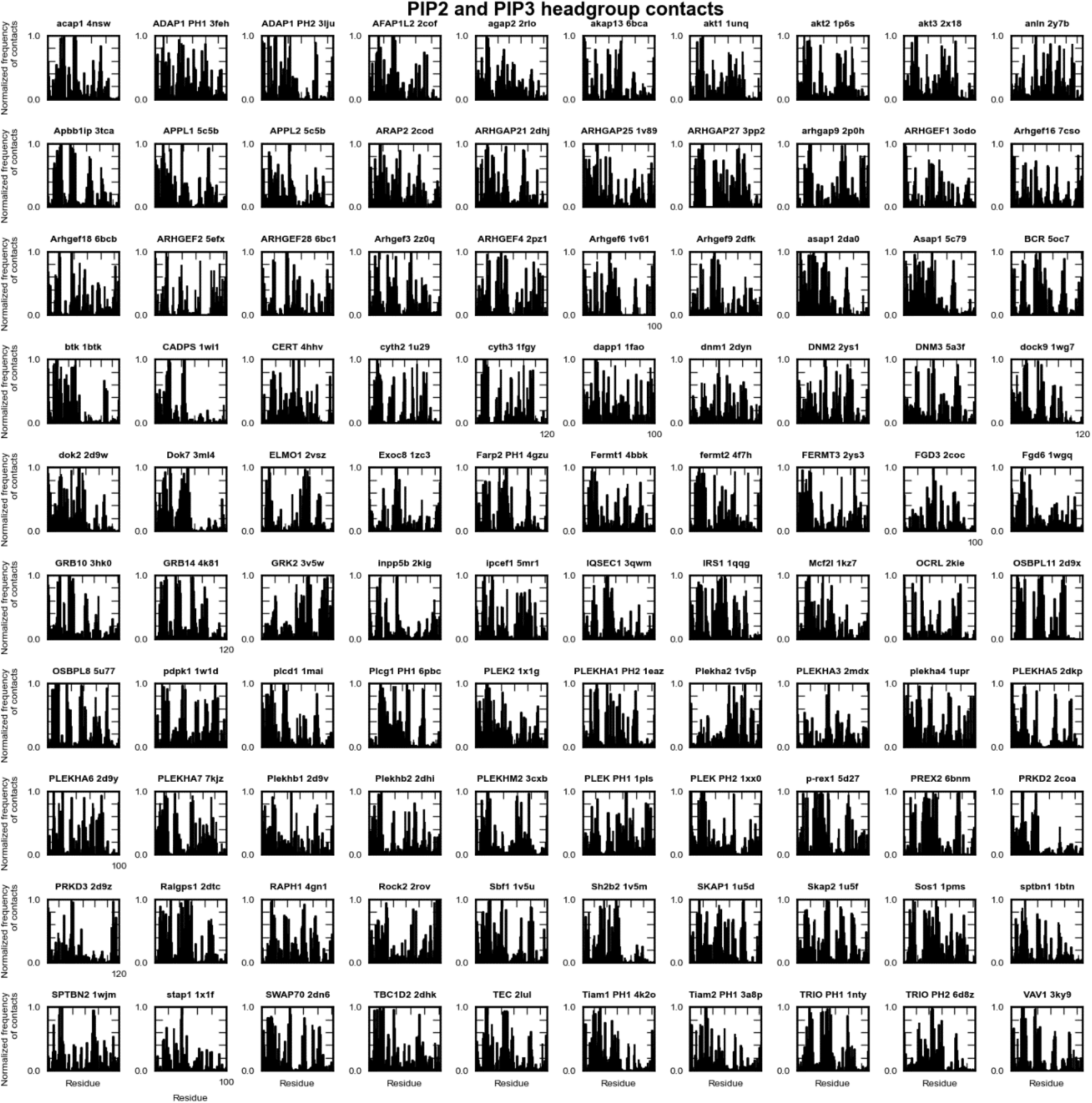
Normalized contacts between PIP_2_/PIP_3_ headgroups and all residues of all PH domains. One contact was counted at a residue for every frame in which a PIP_2_ or PIP_3_ headgroup particle was within a 5.5 Å cut-off distance of any particles of the residue. Contacts were summed over 20 repeat simulations, and then normalized for each PH by dividing the total contacts at each residue by the total contacts made by the residue with the highest number of contacts in that PH domain.

**Table S1.**
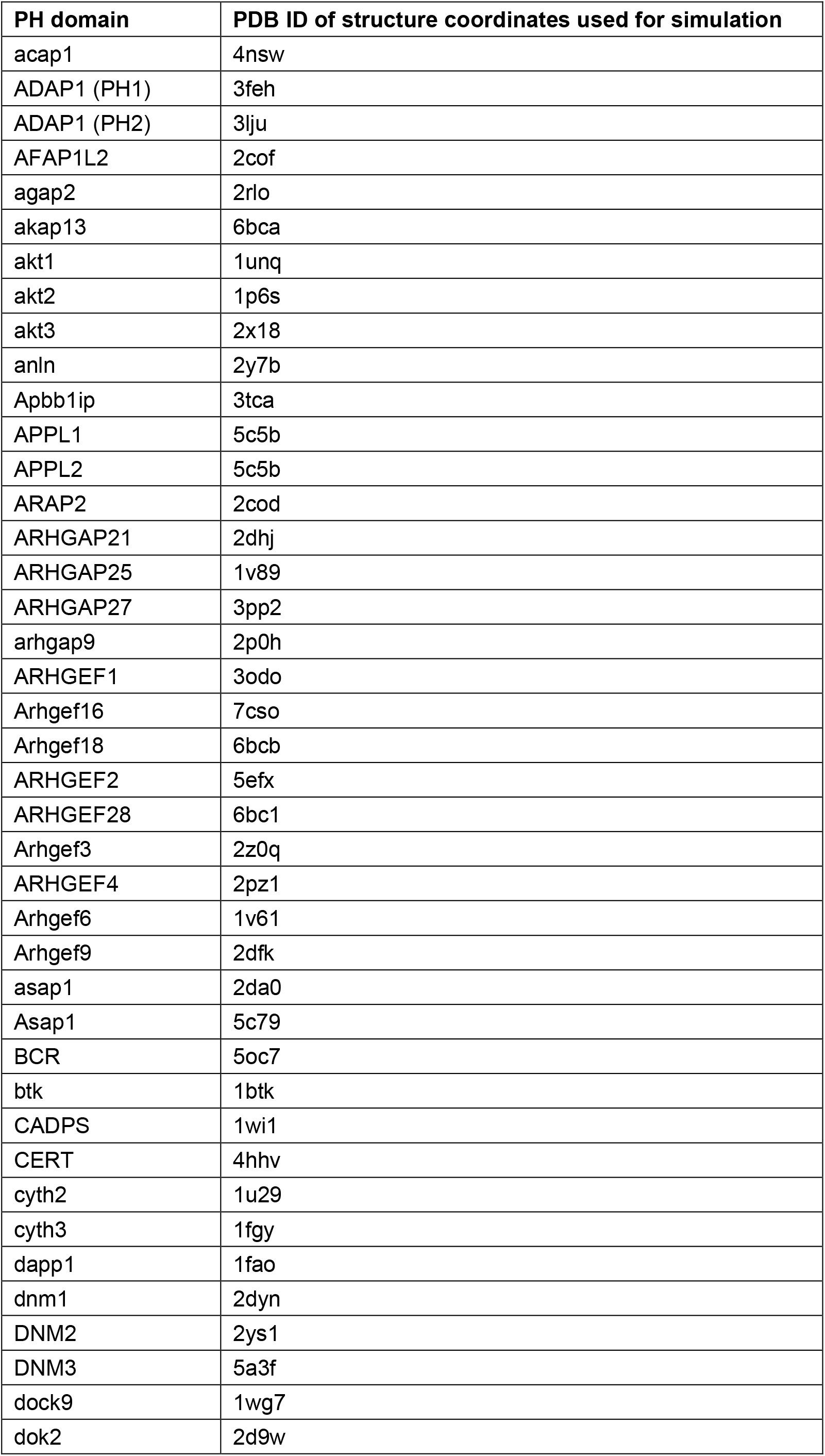
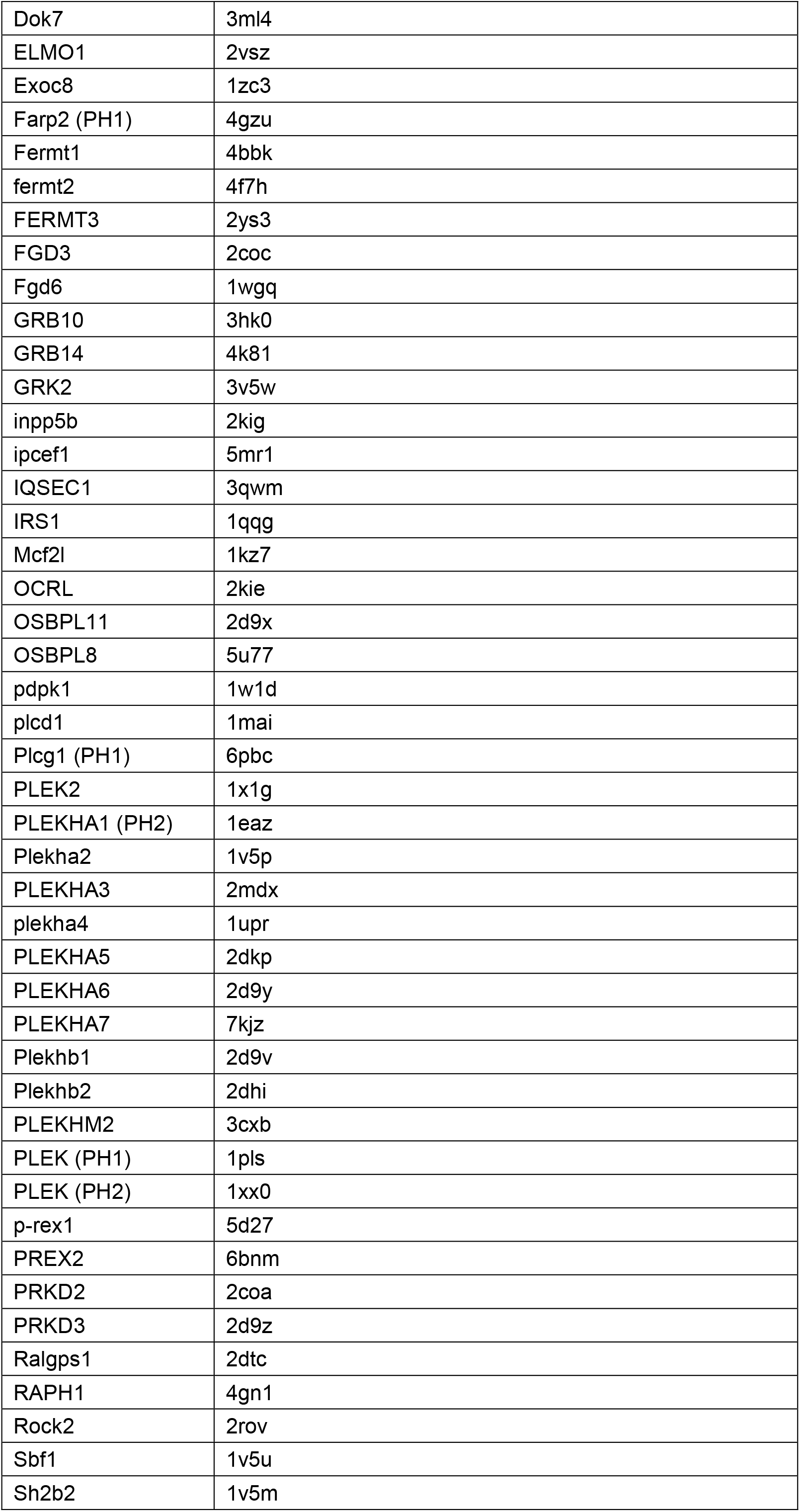
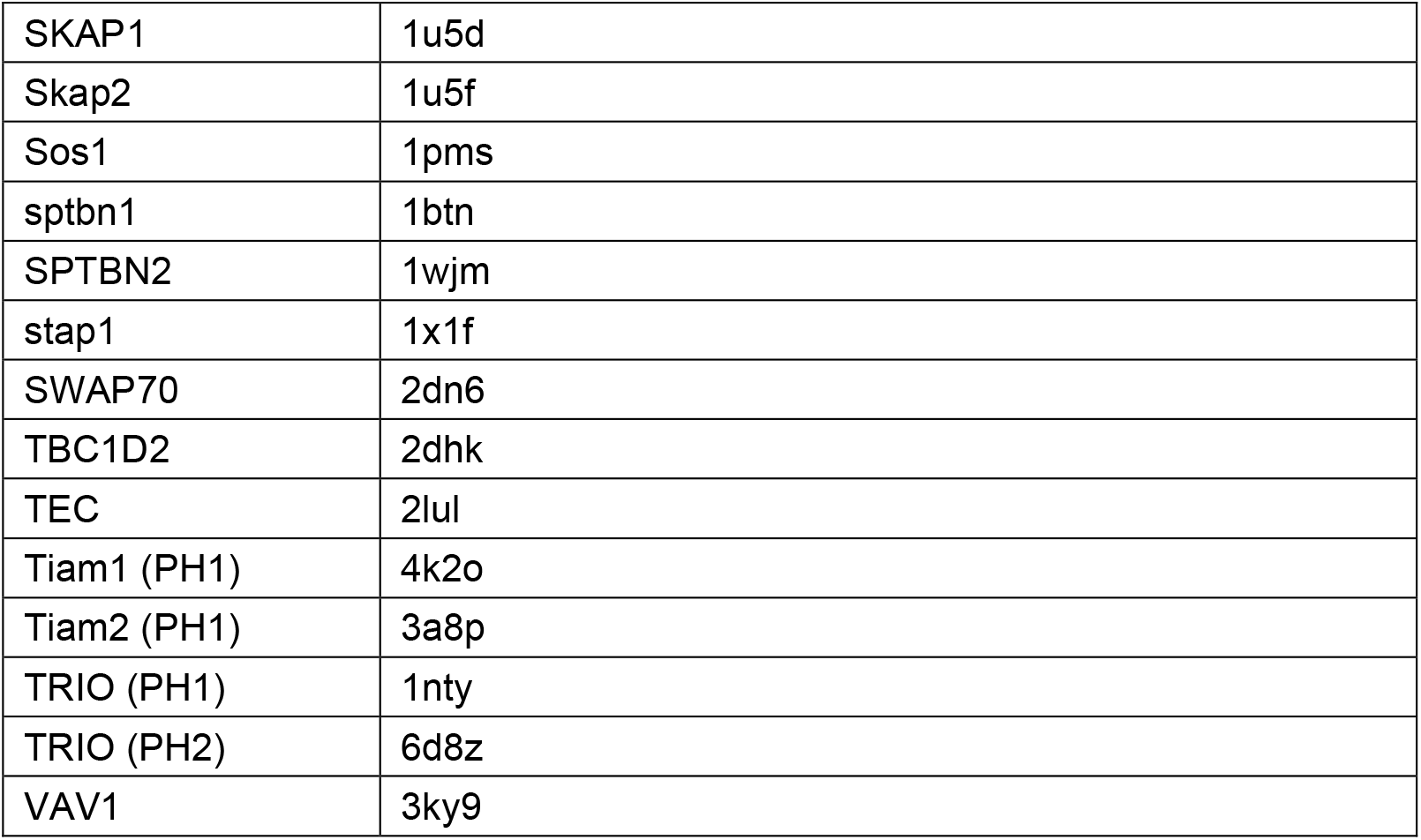
List of all simulated PH domains and the coordinate files used for simulation.

